# Low concordance of short-term and long-term selection responses in experimental *Drosophila* populations

**DOI:** 10.1101/759704

**Authors:** Anna Maria Langmüller, Christian Schlötterer

## Abstract

Experimental evolution is becoming a popular approach to study the genomic selection response of evolving populations. Computer simulation studies suggest that the accuracy of the signature increases with the duration of the experiment. Since some assumptions of the computer simulations may be violated, it is important to scrutinize the influence of the experimental duration with real data. Here, we use a highly replicated Evolve and Resequence study in *Drosophila simulans* to compare the selection targets inferred at different time points. At each time point, approximately the same number of SNPs deviates from neutral expectations, but only 10 % of the selected haplotype blocks identified from the full data set can be detected after 20 generations. Those haplotype blocks that emerge already after 20 generations differ from the others by being strongly selected at the beginning of the experiment and display a more parallel selection response. Consistent with previous computer simulations, our results demonstrate that only Evolve and Resequence experiments with a sufficient number of generations can characterize complex adaptive architectures.

## Introduction

Deciphering the adaptive architecture has been of long-standing interest in evolutionary biology. In contrast to natural populations, experimental evolution (EE) provides the possibility to replicate experiments under controlled, identical conditions and to study how evolution shapes populations in real time (Kawecki et al., 2012). The combination of EE with next-generation sequencing - Evolve and Resequence (E&R) (Long et al., 2015; Schlötterer et al., 2015; Turner et al., 2011) - has become a popular approach to study the genomic response to selection and to identify adaptive loci. E&R has been applied to various selection regimes, such as virus infection (Martins et al., 2014), host-pathogen co-adaptation (Papkou et al., 2019), thermal adaptation (Barghi et al., 2019; Orozco-Terwengel et al., 2012), or body weight (Johansson et al., 2010). A wide range of experimental designs have been used, which vary in census population size, replication level, history of the ancestral populations, selection regime, and number of generations (Burke et al., 2014; Castro et al., 2019; Garland and Rose, 2009; Hardy et al., 2018; Huang et al., 2014; Kawecki et al., 2012; Lang et al., 2013; Michalak et al., 2019; Seabra et al., 2019; Turner et al., 2011). The duration of published E&R studies range from less than 20 generations (Kelly and Hughes, 2018; Rêgo et al., 2019; Turner and Miller, 2012), to a few dozen (Johansson et al., 2010; Orozco-Terwengel et al., 2012) and even hundreds of generations (Burke et al., 2010). Computer simulations showed that the number of generations has a strong influence on the power of E&R studies in sexually reproducing organisms, and increasing the number of generations typically increased it (Baldwin-Brown et al., 2014; Kofler and Schlötterer, 2014; Vlachos and Kofler, 2019). A larger number of generations increases the opportunity for more pronounced allele frequency changes and more recombination events, which results in a more refined mapping resolution (Baldwin-Brown et al., 2014; Kessner and Novembre, 2015; Kofler and Schlötterer, 2014). Since simulations make simplifying assumptions, it is important to scrutinize their conclusions with empirical data. Until recently no suitable data-sets of obligate outcrossing populations were available, which included multiple time points, were replicated, and had starting allele frequencies matching natural populations. We use an E&R experiment (Barghi et al., 2019), which reports allele frequency changes in 10 replicates over 60 generations in 10 generation intervals, to investigate the impact of the experimental duration on the observed genomic response. The time resolved genome-wide polymorphism data of this experiment allow to contrast putative selection targets, which are inferred at different time points, on three analysis levels (candidate SNPs, candidate windows, candidates SNPs shared with reconstructed haplotype blocks, Figure 1). By comparing selection signatures from different time points of the experiment, we show that only a subset of the selection targets are detected at earlier generations, which are not representative of the underlying adaptive architecture.

**Figure 1:**
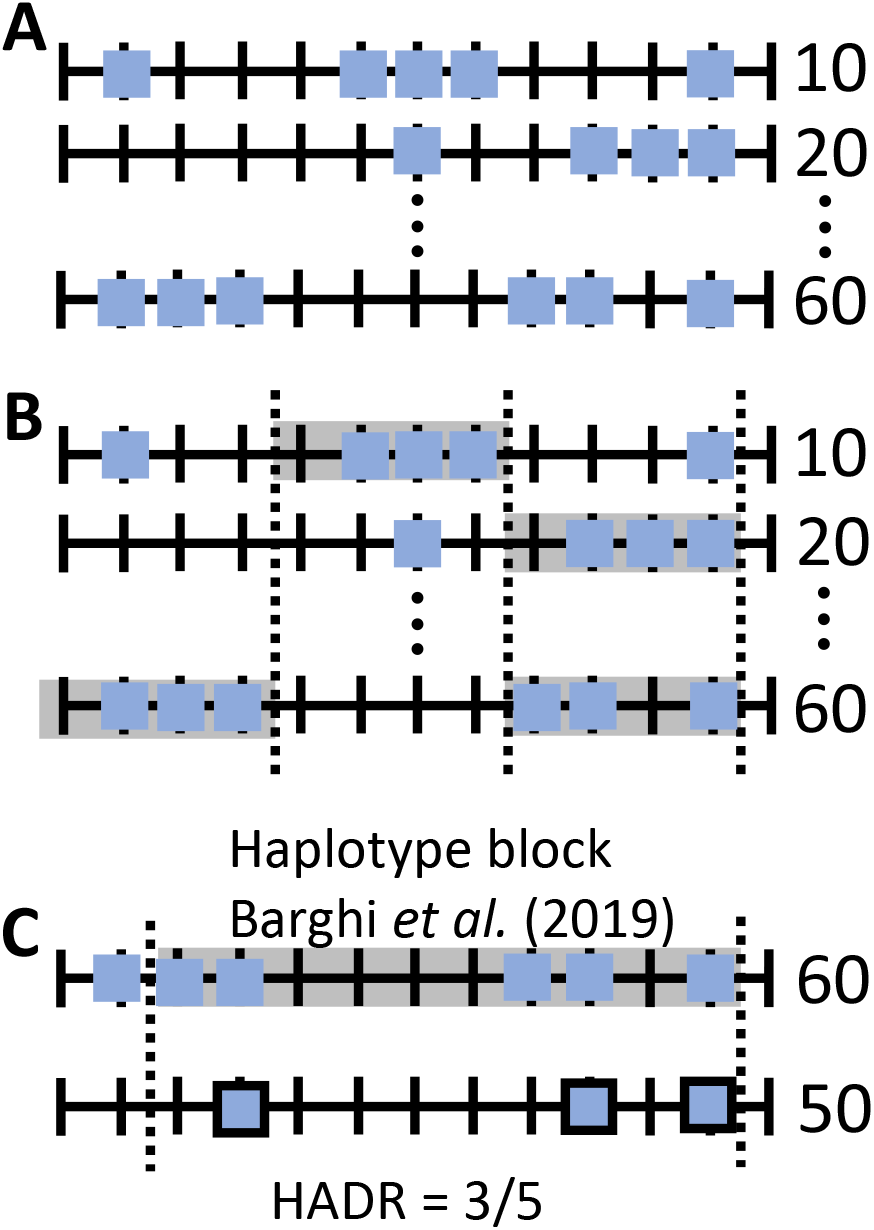
Genomic analysis hierachy: candidate SNPs (**A**), candidate SNPs in a window spanning a fixed number of SNPs (**B**), and candidate SNPs shared with reconstructed haplotype blocks (**C**). (**A**): Candidate SNPs (squares) were determined for each generation and were compared either pairwise (Figure 2), or across multiple time points (Figure 4). (**B**): Window-based approach to detect regions enriched for candidate SNPs. The same number of random SNPs as candidate SNPs were sampled onto the genome repeatedly (n=1 000). Windows (marked by vertical dashed lines) enriched for candidate SNPs contain at least as many candidate SNPs as the 99^*th*^ percentile of the randomly sampled SNPs. Sets of candidate windows were either compared pairwise (Figure 2), or across multiple time points (Figure 4). (**C**): Haplotype block discovery rate (HADR) = the fraction of candidate SNPs (haplotype block marked by vertical dashed lines) that were also discovered at a given time point (framed squares).

## Material & Methods

### Experimental *Drosophila simulans* populations

A detailed description of the *Drosophila simulans* E&R experiment can be found in Barghi et al. (2019) and Hsu et al. (2019). Pooled individuals (Schlötterer et al., 2014) from the evolving populations were sequenced every 10^*th*^ generation starting with the founder population (generation 0) until generation 60 resulting in seven sequenced time points. This E&R experiment started from 202 isofemale lines, which were collected in Tallahassee, Florida. 10 replicate populations evolved in the laboratory at a “cycling hot” temperature regime (12 hours light and 28 °C, 12 hours dark and 18 °C). The census size of the replicates was 1 000 individuals with non-overlapping generations (Barghi et al., 2017, 2019; Hsu et al., 2019).

### Genomic analysis hierarchy

To look for patterns of selection on different scales, we investigated the genomic response of the experimental *Drosophila* populations on three different levels: candidate SNPs, candidate SNPs in a window with a fixed number of SNPs, and candidate SNPs shared with reconstructed selected haplotype blocks. A detailed description for each level is given below, and the different hierarchies are depicted in Figure 1. We performed the same analysis steps at different time points and compared the resulting time point specific selection responses - either pairwise, or across multiple time points - to test the congruence in selection patterns. We evaluated the similarity of two time points with the Jaccard index - a dimensionless parameter ranging from 0 (no overlap) to 1 (sets are identical) - for both candidate SNPs, and candidate windows.

### Identification of candidate SNPs

In the original study, Barghi et al. (2019) applied various filtering steps to obtain a robust SNP set from the ancestral population, resulting in 5 096 200 SNPs on chromosomes X, 2, 3, and 4. A summary of the applied filtering steps can be found in the Supplementary Methods. We used the already determined SNP set of the ancestral population to study the selection response at different time points. For this, we identified “candidate SNPs” based on the frequency difference between the ancestral and evolved populations for each of the six time points (Figure 1A). To identify SNPs with pronounced allele frequency change, we tested (as in Barghi et al. (2019)) replicates separately (Fisher’s exact test) and jointly (Cochran-Mantel-Haenszel test, CMH) using PoPoolation2 (Kofler et al., 2011). We chose the CMH test statistic out of a range of approaches that allow to identify selected SNPs, because it has been shown recently that the CMH test – although not taking intermediate time points into account – consistently outperforms other methods regardless of the investigated selection scenario (Vlachos et al., 2019). As outlier SNPs with extreme read depths had already been removed in the original study, additional minimum and maximum read depth restrictions were not imposed on the SNP set. Neither of the two chosen test statistics to determine candidate SNPs (CMH test, Fisher’s exact test) account for allele frequency change due to genetic drift. To detect SNPs that show more allele frequency change than expected under drift, we performed neutral simulations with Nest (Jónás et al., 2016) using estimates of the effective population size (*N_e_*) between generation 0 and the focal time point (Table S1-S3). The neutral simulations further used the empirical starting allele frequencies and read depths. For the CMH test, *N_e_* estimates were averaged across replicates for autosomes, and the X chromosome separately. For Fisher’s exact test, we used replicate-specific *N_e_* estimates of the autosomes. Based on these neutral simulations we determined candidate SNPs with a false discovery rate smaller than 5 % (Barghi et al., 2019).

We identified 56 166 candidate SNPs in generation 60, compared to 55 199 in Barghi et al. (2019). This small discrepancy can be explained by stochastic differences arising from the neutral simulations used to determine the significance threshold. We excluded six out of 99 haplotype blocks from Barghi et al. (2019) with less than 90% of the previously reported candidate SNPs.

### Identification of candidate windows

Because of linkage in the experimental populations (Nuzhdin and Turner, 2013; Tobler et al., 2014), the first genomic analysis level - the number of candidate SNPs (Figure 1A) - will most likely suffer from an excess of candidate loci. We used a window-based approach (Figure 1B) as second genomic analysis hierarchy to account for a potential excess of candidate SNPs. We split the main chromosomes (X, 2 and 3) into non-overlapping windows of 5 000 SNPs that are segregating in all generations and replicates. We chose SNPs instead of base pairs as window size measure in order to allow for variation in SNP density along the genome. To determine if a given window displays a potential selection response(= it contains more candidate SNPs than expected), we sampled the same number of random SNPs as candidate SNPs in this window (1 000 iterations) onto the whole genome. “Candidate windows” contained at least as many candidate SNPs as the 99^*th*^ percentile of randomly sampled SNPs. We received time point specific candidate window sets (Figure 1B, S1), by applying the procedure described above independently to candidate SNPs from all time points. To check for the robustness of candidate window patterns, we varied the window size, and allowed SNPs to fix during the experiment, which both resulted in qualitatively similar results (Table S4, and S5).

Candidate windows (=the number of candidate SNPs in a window) is a summary statistic which ignores the significance of the candidate SNPs. If a signal is robust between two time points, we expect the same *p*-value-based ranking of candidate SNPs assuming homogeneous read depth for all sites and time points, and that candidate SNPs do not fix during the experiment. Although read depth heterogeneity among sites will change the confidence in the estimates of the allele frequency change, the average dynamics of relative significance allow us to determine whether the robustness of a putative selection response increases with time. Thus, we also evaluated whether candidate SNPs in a given window had a similar significance rank. For each candidate window we created a ROC-like curve (similar to Jakšić and Schlötterer (2016)) by ranking the candidate SNPs by their *p*-values - the most significant SNP was assigned rank 1 - and calculating the overlap in top-ranked SNPs between two time points.

### HAplotype block Discovery Rate (HADR)

The third genomic analysis level is based on reconstructed selected haplotype blocks of the original study (Figure 1C): Barghi et al. (2019) clustered candidate SNPs from generation 60 into selected haplotype blocks based on similar allele frequency trajectories over time and replicates (Franssen et al., 2016) to assess the underlying haplotype structure in the experimental populations. The reconstructed haplotype blocks were further validated with experimentally phased haplotypes from ancestral and evolved populations (Barghi et al., 2019), and 96 % of the reconstructed haplotype blocks could be confirmed by the experimentally derived haplotypes. This suggests that reconstructed haplotype blocks provide a robust set of linked candidate SNPs that can be used to investigate time point specific patterns of selection.

Taking advantage of this additional confirmation of the candidate SNPs in a selected haplotype block, we developed a third measure of similarity between time points. We determined the fraction of candidate SNPs comprising a haplotype block that were also discovered at a given time point (haplotype block discovery rate, HADR, Figure 1C) using the `poolSeq` R-package (Taus et al., 2017). We note that the inference of selected haplotype blocks at each generation does not provide a good alternative to the HADR measure. It has been shown that the ability of the clustering method to group SNPs into haplotype blocks is dependent on the number of time points (Franssen et al., 2016). This results in less power at early generations where fewer time points are available.

### Early Detected HAplotype blocks (EDHAs)

We applied various clustering methods (hierarchical clustering (Pollard and Laan, 2005), PCA, and kmeans (Hartigan and Wong, 1979)) to group reconstructed haplotype blocks by their HADR patterns over time. The hyper-parameter k (which determines the number of clusters) in the kmeans clustering procedure was set to 5 based on the gap statistic (Tibshirani et al., 2001). Both the k-means clustering and the PCA revealed a clearly distinguishable group of 10 haplotype blocks with elevated HADR early on in the experiment (Figure S2, S3). We refer to the haplotype blocks in this cluster as early detected haplotype blocks (EDHAs). We investigated whether EDHAs have distinct characteristics that might explain why they are early detectable by comparing the following features between EDHAs, and non-EDHAs: haplotype block length, median starting allele frequency, average recombination rate (Howie et al., 2019), selection coefficient (*s*, estimated with `poolSeq` (Taus et al., 2017)) in generation 20, *s* in generation 60, selection coefficient ratio 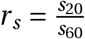, and number of rising replicates in generation 20 and in generation 60. To avoid that non–responding replicates bias the selection coefficient estimates downwards, we averaged selection coefficients over replicates that passed a certain allele frequency change threshold. Following Barghi et al. (2019), we classified a haplotype block as replicate specific, if the allele frequency of candidate SNPs from a haplotype block increases on average by at least 10%. To test the robustness of the replicate specific rising behaviour, we also used 5, 15, and 20% thresholds, which all resulted in qualitatively similar results.

### Simulation of an E&R experiment

#### Simulation with linkage

Despite the selected haplotype blocks were inferred with high confidence by Barghi et al. (2019), we were interested to confirm our conclusions with a simulated data set for which the selection targets are known. Because the distinction between true causative SNPs and linked polymorphisms is challenging (Nuzhdin and Turner, 2013; Tobler et al., 2014), we obtained a simulated data set that models linkage and resembles the Barghi et al. (2019) study using MimicrEE2 (v. 206) (Vlachos and Kofler, 2018). Our ancestral population was built from 189 ancestral haplotypes published in Barghi et al. (2019) and Howie et al. (2019). We simulated a selective sweep scenario with 99 independent selection targets, where each of them is located in the inferred selected haplotype block, has the same starting frequency, and selection coefficient (both estimated as the median from all selected SNPs in the block) (Barghi et al., 2019). Since we were primarily interested in the early genomic responses after the populations has been exposed to a new environment, we reasoned that selective sweeps approximate the selection trajectories for a polygenic trait with stabilizing selection (as presented in Barghi et al. (2019)) rather well (Chevin and Hospital, 2008; Höllinger et al., 2019; Jain and Stephan, 2017a,b). Using the *Drosophila simulans* recombination rate map (Howie et al., 2019), we simulated 10 outcrossing populations with constant census size adapting for 60 non-overlapping generations. We sampled the read depth for each site from a Poisson distribution with *λ* equal to the average read depth for each population and time point as reported in Barghi et al. (2019). For each site, we applied binomial sampling with the determined read depth and true allele frequency to mimic the sampling of reads out of a DNA pool (Jónás et al., 2016). This forward simulation results in a data set very similar to the original study that allows us to analyze selection patterns on all three genomic analysis hierarchy levels, but in contrast to the empirical data, the causative SNPs are known. The identification and analysis of candidate SNPs and windows of the simulation followed the protocol of the empirical data (as described above; Figure 1AB). Instead of relying on the dynamics of selected haplotype blocks (see HADR, Figure 1C), we investigated the number of selection targets that can be found amongst statistical outliers across generations. Early detected targets (see EDHAs) were defined as targets that are consistently detected as outlier from generation 20 onwards.

#### Simulation without linkage

To explicitly test whether the highly parallel selection signature of EDHAs at generation 20 can be explained by synergistic effects of drift and selection, we simulated 100 000 co-dominant, unlinked loci in 10 replicate populations with equal starting allele frequency (10 %, median starting allele frequency reported in Barghi et al. (2019)), and selection coefficient (s=0.05, median selection coefficient reported in Barghi et al. (2019)) using the `poolSeq` R-package (Taus et al., 2017). Based on neutral simulations (conducted with the the `poolSeq` R-package (Taus et al., 2017), we determined candidate loci with a false discovery rate smaller than 5 % after 20 generations (CMH test) (Barghi et al., 2019), and compared the parallelism of candidate loci to the parallelism of non-candidate loci. If the synergistic effect of drift and selection is causing elevated parallelism, we would expect candidate loci to rise in significantly more replicates than their non-significant counterparts, despite having the same starting allele frequency and selection coefficient.

## Results & Discussion

### Subsequent time points are more similar for advanced generations

We studied the similarity of selection signatures for different time points using 10 replicates of a *D. simulans* population, which evolved for 60 generations to a novel hot environment (Barghi et al., 2019). With Pool-Seq (Schlötterer et al., 2014) data from every 10^*th*^ generation, we evaluated the selection signature on three different levels: candidate SNPs, candidate SNPs in a window spanning a fixed number of SNPs, and candidate SNPs shared with reconstructed selected haplotype blocks (Figure 1). The similarity of inference between two time points was determined by the Jaccard index (J), ranging from 0 (no overlap between two SNP sets) to 1 (sets are identical). We found that all candidate SNP sets are more similar than expected by chance. The Jaccard index ranged from 0.08 (generation 10 vs generation 60) to 0.40 (generation 50 vs generation 60), where subsequent time points are more similar than those separated for more than 10 generations (e.g. J=0.15 (generation 10 vs generation 20); J=0.08 (generation 10 vs generation 60)). Furthermore, the similarity of candidate SNP sets from subsequent time points increased with time until it ultimately more than doubles for the last two generations (J=0.15 (generation 10 vs generation 20); J=0.34 (generation 50 vs generation 60), Figure 2A). The monotonic increase in similarity with time shows that more similar selection patterns are detected at later generations. While this suggests that outliers at generation 60 provide the most reliable selection signature, we were interested to confirm this with computer simulations where all selection targets and the type of selection are known. We simulated an E&R experiment that resembles the data from Barghi et al. (2019) and repeated our analysis on simulated, time resolved allele frequencies. Again, we observed that candidate SNP sets are more similar than expected by chance, the majority of subsequent time points are more similar than those separated for more than 10 generations, and the similarity of candidate SNP sets from subsequent time points increases with time (Figure S4).

**Figure 2:**
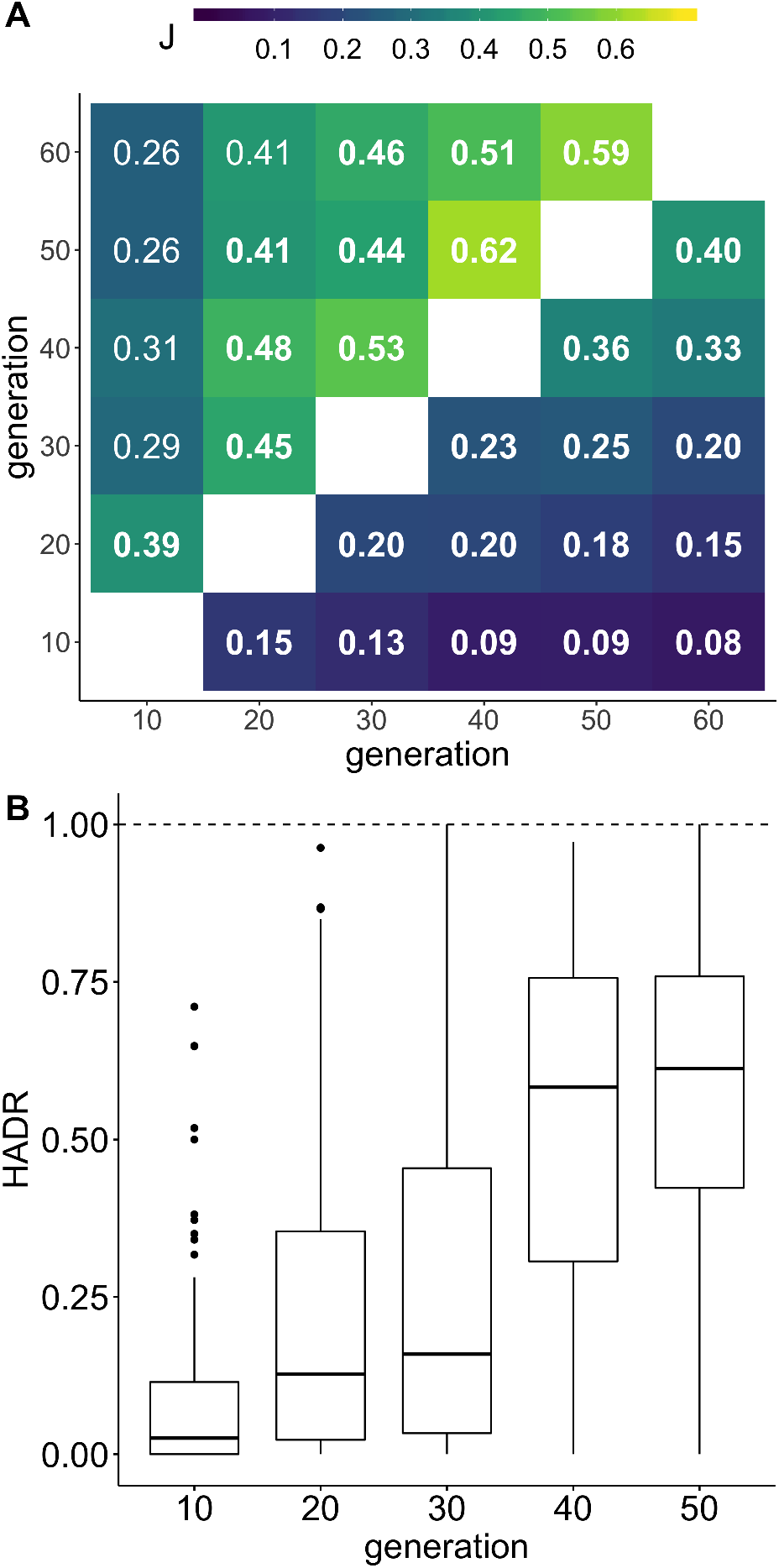
Similarity measures for candidate SNPs, candidate SNPs in a window with a fixed number of SNPs, and candidate SNPs shared with reconstructed selected haplotype blocks (**A**): Jaccard index (J) for pairwise comparisons of candidate sets. The top triangle shows candidate window sets, the bottom triangle candidate SNP sets. Significant similarities (*p*-value <0.05 after multiple testing correction, 10 000 bootstraps) are written in bold.(**B**): The rate at which selected SNPs of 93 haplotype blocks from generation 60 were already discovered at earlier generations (haplotype discovery rate, HADR).

Since the analysis of single SNPs suffers from considerable stochasticity, and neighboring SNPs are not independent (Howie et al., 2019; Tobler et al., 2014), we repeated the analysis of different time points using non-overlapping windows of 5 000 SNPs (Figure 1B). Reasoning that windows containing a target of selection will harbor multiple candidate SNPs, we defined selected windows as those, which harbor more candidate SNPs than expected by chance. Consistent with higher stochasticity at the SNP level, a higher similarity was observed for candidate windows (from J=0.26 (generation 10 vs generation 60) to J=0.62 (generation 40 vs generation 50)). Again, adjacent time points have a higher Jaccard index than time points farther apart (J=0.26 (generation 10 vs generation 60); J=0.39 (generation 10 vs generation 20)). The similarity of subsequent time points also increases with the duration of the experiment (J=0.39 (generation 10 vs generation 20); J=0.59 (generation 50 vs generation 60), Figure 2A). In contrast to the SNP level, the set of selected windows after 10 generations is only significantly similar to generation 20, but not to any other generation. Thus, the pattern of reduced similarity of selection targets in the early generations is confirmed at the window level, albeit with different significance levels. The same pattern was noticed for the simulated E&R data (Figure S4).

For an alternative measure of similarity, we used the ranking of candidate SNPs in a specific window based on their *p*-values and compared it between different time points. If a signal is robust between two time points, we expect the same SNP ranking of segregating SNPs in a selected window. Consistent with the other tests, we found that the congruence in candidate SNP ranking increases with time in both empirical and simulated data (Figure 3, S5). To rule out that rare SNPs are responsible for the dissimilarity between early and late time points, we calculated similarity measures based on SNPs that are segregating at all generations and time points. Nevertheless, including SNPs which were lost in at least one replicate during the experiment did not result in a pronounced decrease in similarity (Figure S6).

**Figure 3:**
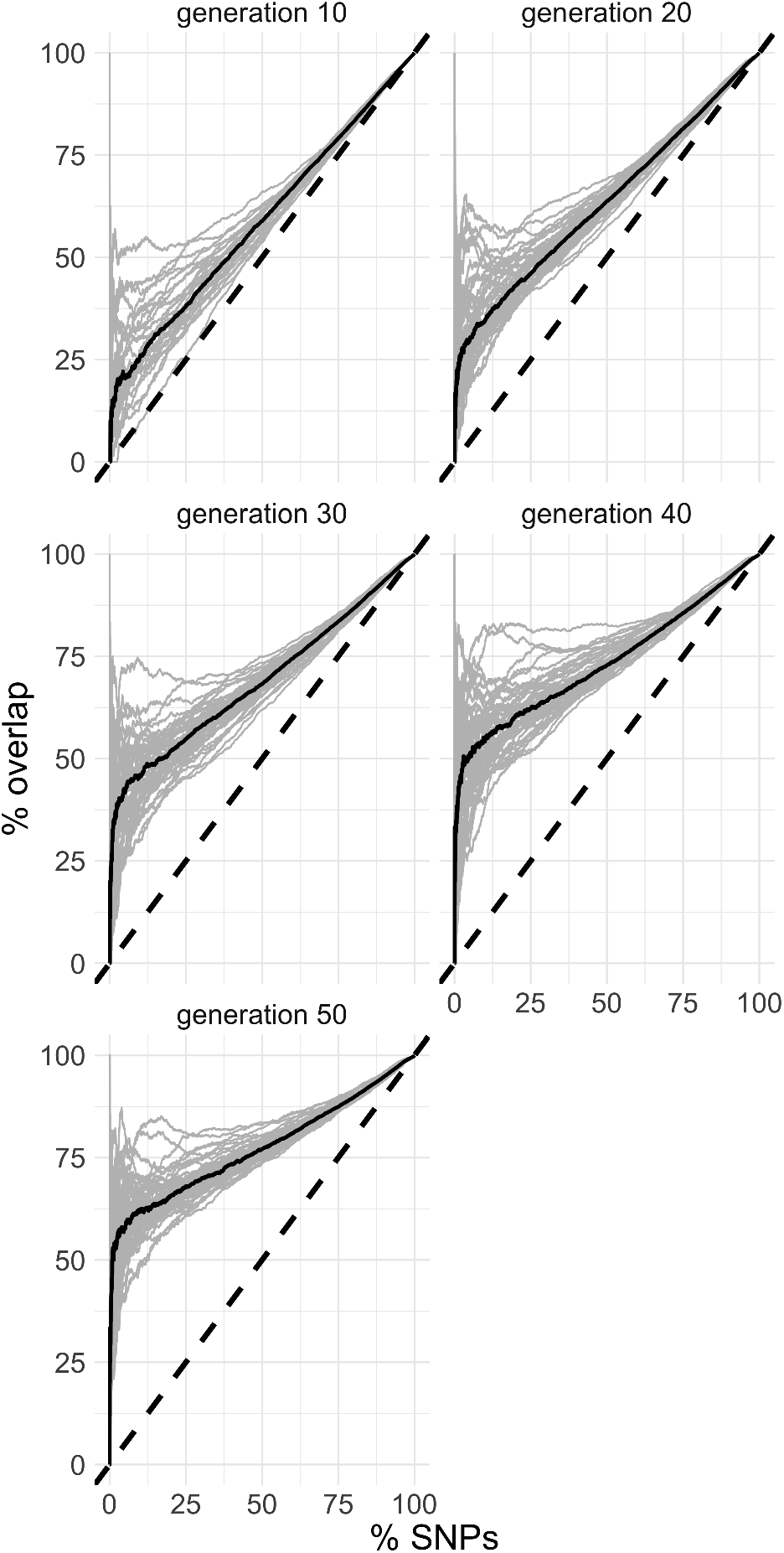
The rank of candidate SNPs becomes more congruent with time. In this ROC-like graph, the ranking of all candidate SNPs in candidate windows is compared. Each panel shows one intermediate time point compared to generation 60. The overlap (in percent) for each candidate window is indicated by a separate line. The median overlap (solid black line) monotonically increases with experimental duration, demonstrating that the ranking of candidate SNPs is more robust for advanced generations. The black, dashed lines shows the expected overlap in SNP ranking if every variant at generation 60 is recapitulated in a previous time point.

The analysis of selected haplotype blocks provides another possibility to control for non-independence of single candidate SNPs. We calculated the haplotype block discovery rate (HADR, Figure 1C) – the fraction of candidate SNPs in a haplotype block that are rediscovered at a given time point. Similar to the other analyses, we observe higher similarity between later time points (Figure 2B), with a pronounced increase of median HADR between generation 30 (< 25 %) and 40 (> 50 %). In the simulated data, the number of detected targets monotonically increased until generation 50 (Figure S7).

Independent of the measure of similarity between time points, for both empirical and simulated data, the selection signatures of early time points are consistently less similar than those from later time points. Under the assumption that the same alleles are under positive selection over the whole time span of the experiment, this observation highlights that a more reliable identification of selection targets requires longer experiments with additional generations. With the limitation that for empirical data the true targets of selection are not known, in the Barghi et al. (2019) data a particularly striking observation is that a similar number of candidate SNPs was detected at each time point despite the number of “true” targets increased with the duration of the experiment (Table S6). While this suggests that earlier time points harbor more false positives, we would like to point out that the low concordance in selection patterns between early, and late generations in the experiment could also be caused by a change (or a even a reversal) in selection. It may be possible that some targets were only selected during the first generations, and that the strength of selection changed later on. This may be caused for example either by epistatic interactions, or an unobserved change in our experimental environment. While this cannot be ruled out for the empirical data, this cannot explain the similarity dynamics in the sweep simulation with additive selection, and no epistasis modeled (Figure S4, S7).

It is important to note that the sweep simulation did not have a similar number of candidate SNPs over time, rather the number of false positives increased with the duration of the experiment (Table S7). We attribute this discrepancy of the simulated and empirical data outcome to the selection regime employed, which affects mainly later generations. Sweep simulations result in haplotype blocks with many selection targets (Barghi and Schlötterer, 2020), which in turn influences the number of candidate SNPs. Simulations of polygenic adaptation can become quite complex and rich in parameters (Thornton, 2019), which limits the power of computer simulations to scrutinize this result further. Rather, experimental validation of selection targets in secondary E&R studies (Burny et al., 2020) may be used to confirm selection signatures beyond statistical testing and could serve an important role to test early selection signatures that cannot be confirmed at later time points.

### Only few selection targets are shared across all generations

More than 27 000 candidate SNPs can be identified at each time point (Table S6), but only a small (5 %) subset is consistently detected at every generation (Figure 4; including rare SNPs see Figure S8). The small subset of consistently detected candidate SNPs can not be explained by fixation of candidate SNPs of early generations before generation 60 - the ratio of fixed candidate SNPs does not exceed 3.5 % at any consecutive time point (Figure S9). Apart from highlighting the more robust selection signatures with an increasing number of generations, this analysis raises an important concern about the usefulness of meta-analyses on the SNP level. With less than 5% of the SNPs being shared in the same selection experiment, it will be extremely difficult to compare studies that started from different founder populations and were selected for a different number of generations.

**Figure 4:**
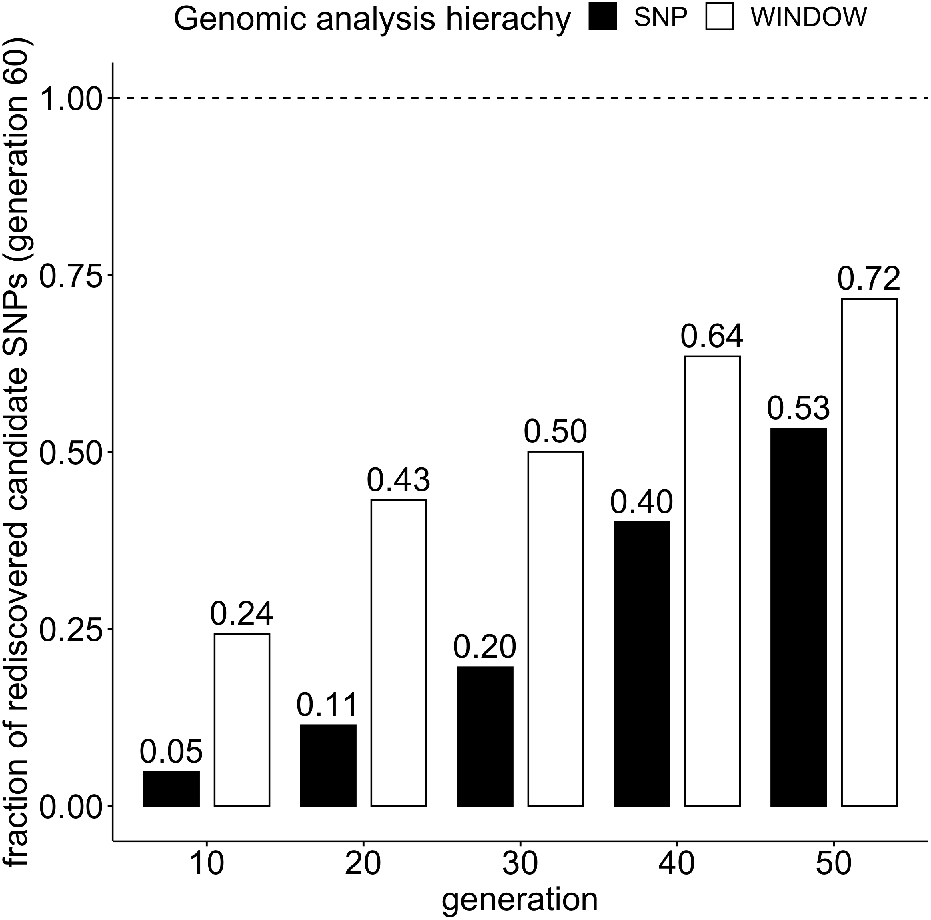
5% of candidate SNPs in generation 60 are detected consistently at every generation. The bars depict the fraction candidate SNPs (black) and candidate windows (white) at generation 60, which are candidates in all subsequent generations (e.g. 40% of generation 60 candidate SNPs are candidates in generation 50 and 40). Candidate windows are more consistent than candidate SNPs. Figure S8 depicts the ratios for candidate sets that are not restricted to SNPs segregating in all generations and time points.

We repeated the analysis for windows and determined the number of selected windows that are shared across all generations. With 18 out of 74 candidate windows in generation 60 (24.3 %, Figure 4) being detected at all generations, the window analysis shows more consistency across time points than a SNP-based analysis. This observation is independent of window size (Table S4, and S5), and the inclusion of rare SNPs into the analysis (Figure S8). Because we observed a similar trend in our simulated experiment (Figure S10), we propose that meta-analyses of E&R data should be performed on the level of windows, or based on selected haplotype blocks to avoid false negatives due to the high stochasticity of SNP-based analyses.

### Selection signatures detected early in the experiment are not representative of the underlying adaptive architecture

This study focused on the comparison of selection targets detected at early and late time points. Since analyses based on single SNPs are very stochastic, we investigated the fraction of candidate SNPs comprising a haplotype block that were also discovered at earlier time points (HADR, Figure 1C). We detected 10 haplotype blocks with elevated HADR in generation 20 (early detected haplotype blocks, EDHAs, Figure S2, S3). We found that EDHAs do not differ in their starting allele frequency, haplotype block length, average recombination rate, absolute selection coefficients, or number of rising replicates after 60 generations from other haplotype blocks (Figure S11). EDHAs are, however, more strongly selected at the beginning of the experiment, but are equally strongly selected as the remaining haplotype blocks at later generations (Figure 5A). Consistent with stronger selection at earlier time points, the selection signature of EDHAs is significantly more parallel across replicates after 20 generations of adaptation in both empirical and simulated data. (Figure 5B, S12). We attribute this observation to a phenomenon similar to the “winners curse”, that is that loci where stochastic effects increased the frequency in multiple replicates to enhance the contribution by selection are more likely to be detected. We explicitly tested this interpretation with computer simulations. For this, we simulated unlinked loci with identical starting frequencies and selection coefficients. As expected not all selected loci are detected after 20 generations, but detected loci show a more parallel selection signature than not detected ones despite having the same starting allele frequency, and selection coefficient (Figure S13).

**Figure 5:**
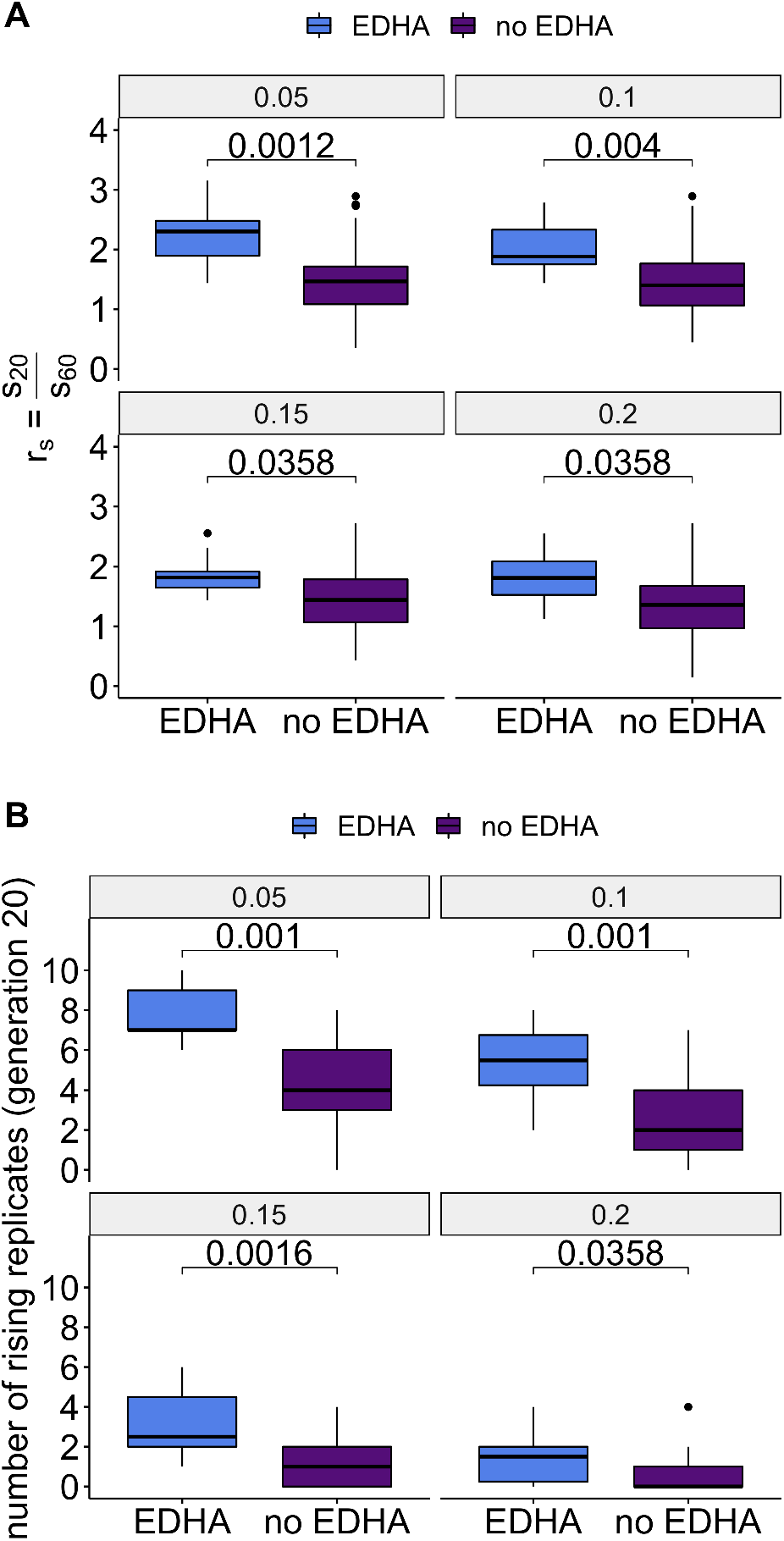
Early Detectable HAplotype blocks (EDHAs) differ from the other selected haplotype blocks. (**A**): The ratio of selection (*r*_*s*_) coefficients determined for early generations (generation 20, *s*_20_) and late generations (generation 60, *s*_60_) is significantly higher for EDHAs. (**B**): EDHAs rise in more replicates than other haplotype blocks after 20 generations. Both observations are robust to different allele frequency change thresholds. Values above the boxplots represent the two-tailed Mann-Whitney test *p*-values corrected for multiple testing with the Benjamini-Hochberg procedure.

All statistical tests, which are evaluating a parallel selection signature across replicates, are more likely to detect selection signatures shared across replicates, even with only moderate allele frequency changes. This “enhanced” parallelism could result in wrong or incomplete conclusions about the underlying genetic architecture (Jain and Stephan, 2017a; Höllinger et al., 2019; Thornton, 2019). The analysis of selection signatures in replicated experiments running for only a moderate number of generations is more likely to detect parallel than replicate specific selection signatures. This bias is not restricted to our study, but also an experimental study of *D. simulans* populations adapting 10 to 20 generations to a new temperature regime (Kelly and Hughes, 2018) found more parallel selection responses. We propose that additional analyses contrasting selection signatures of early and late time points are needed to confirm the enrichment of parallel selection signatures in short-term experiments.

## Acknowledgments

We thank the members of the Institut für Populationsgenetik for fruitful discussion and support. Special thanks to Neda Barghi, Sheng-Kai Hsu and Claire Burny for helpful comments on earlier versions of the manuscript. This work was supported by the Austrian Science Fund (FWF, grant number W1225) and an European Research Council (ERC) grant (ArchAdapt).

## Data Accessibility

- Data of the original study: Barghi et al. (2019), Data from: Genetic redundancy fuels polygenic adaptation in Drosophila, Dryad, Dataset, https://doi.org/10.5061/dryad.rr137kn For this study, we used the F0–F60 sync file, and the haplotype block assignment of sin-gle SNPs (file : F0-F60SNP_CMH_FET_blockID.sync.zip); the estimated selection coefficients, and starting allele frequencies of reconstructed haplotype blocks (file: S_SAF_0.1AFC.txt), as well as the characteristics of the reconstructed haplotype blocks (https://doi.org/10.1371/journal.pbio.3000128.s014) and the coverage of SNPs for all time points and replicates https://doi.org/10.1371/journal.pbio.3000128.s021)
- Phased ancestral haplotypes and recombination map: Howie et al. (2019), Data from: DNA motifs are not general predictors of recombination in two Drosophila sister species., Dryad, Dataset, https://doi.org/10.5061/dryad.744p394 For this study we used the file Dsim_recombination_map_LOESS_100kb_1.txt (recombination map for the simulations), and the phased *D. simulans* haplotypes.
- All the necessary scripts for reproducing the simulations and results are available on GitHub (https://github.com/AnnaMariaL/LowConcordance_ER).

## Authors Contribution

CS designed the experiment. CS and AML designed the analysis. AML performed the bioinformatic analysis. CS and AML wrote the manuscript.

## Notes

### Competing Interest Statement

The authors have declared no competing interest.

### Summary of Updates

- additional simulations to support hypothesis - work on figures

## References

Baldwin-Brown, J. G., Long, A. D., and Thornton, K. R. (2014). The power to detect quantitative trait loci using resequenced, experimentally evolved populations of diploid, sexual organisms. Molecular Biology and Evolution, 31(4):1040–1055.

Barghi, N. and Schlötterer, C. (2020). Distinct Patterns of Selective Sweep and Polygenic Adaptation in Evolve and Resequence Studies. Genome Biology and Evolution, 12(6):890–904.

Barghi, N., Tobler, R., Nolte, V., Jakšić, A. M., Mallard, F., Otte, K. A., Dolezal, M., Taus, T., Kofler, R., and Schlötterer, C. (2019). Genetic redundancy fuels polygenic adaptation in *Drosophila*. PLOS Biology, 17(2):1–31.

Barghi, N., Tobler, R., Nolte, V., and Schlötterer, C. (2017). *Drosophila simulans*: A species with improved resolution in evolve and resequence studies. G3: Genes, Genomes, Genetics, 7(7):2337–2343.

Burke, M. K., Dunham, J. P., Shahrestani, P., Thornton, K. R., Rose, M. R., and Long, A. D. (2010). Genome-wide analysis of a long-term evolution experiment with *Drosophila*. Nature, 467(7315):587–590.

Burke, M. K., Liti, G., and Long, A. D. (2014). Standing genetic variation drives repeatable experimental evolution in outcrossing populations of *Saccharomyces cerevisiae*. Molecular Biology and Evolution, 31(12):3228–3239.

Burny, C., Nolte, V., Nouhaud, P., Dolezal, M., Schlötterer, C., and Baer, C. (2020). Secondary Evolve and Resequencing: An Experimental Confirmation of Putative Selection Targets without Phenotyping. Genome Biology and Evolution, 12(3):151–159.

Castro, J. P., Yancoskie, M. N., Marchini, M., Belohlavy, S., Hiramatsu, L., Kucčka, M., Beluch, W. H., Naumann, R., Skuplik, I., Cobb, J., Barton, N. H., Rolian, C., and Chan, Y. F. (2019). An integrative genomic analysis of the longshanks selection experiment for longer limbs in mice. eLife, 8:e42014.

Chevin, L. M. and Hospital, F. (2008). Selective sweep at a quantitative trait locus in the presence of background genetic variation. Genetics, 180(3):1645–1660.

Franssen, S. U., Schlötterer, C., and Barton, N. H. (2016). Reconstruction of Haplotype-Blocks Selected during Experimental Evolution. Molecular Biology and Evolution, 34(1):174–184.

Garland, T. and Rose, M. (2009). Experimental Evolution: Concepts, Methods, and Applications of Selection Experiments. University of California Press.

Hardy, C. M., Burke, M. K., Everett, L. J., Han, M. V., Lantz, K. M., and Gibbs, A. G. (2018). Genome-Wide Analysis of Starvation-Selected *Drosophila melanogaster*-A Genetic Model of Obesity. Molecular Biology and Evolution, 35(1):50–65.

Hartigan, J. A. and Wong, M. A. (1979). Algorithm AS 136: A K-Means Clustering Algorithm. Journal of the Royal Statistical Society. Series C (Applied Statistics), 28(1):100–108.

Höllinger, I., Pennings, P. S., and Hermisson, J. (2019). Polygenic adaptation: From sweeps to subtle frequency shifts. PLOS Genetics, 15(3):1–26.

Howie, J. M., Mazzucco, R., Taus, T., Nolte, V., and Schlötterer, C. (2019). DNA Motifs Are Not General Predictors of Recombination in Two *Drosophila* Sister Species. Genome Biology and Evolution, 11(4):1345–1357.

Hsu, S.-K., Jaksic, A. M., Nolte, V., Barghi, N., Mallard, F., Otte, K. A., and Christian, S. (2019). A 24 h Age Difference Causes Twice as Much Gene Expression Divergence as 100 Generations of Adaptation to a Novel Environment. Genes, 10(2):89.

Huang, Y., Wright, S. I., and Agrawal, A. F. (2014). Genome-wide patterns of genetic variation within and among alternative selective regimes. PLOS Genetics, 10(8):1–15.

Jain, K. and Stephan, W. (2017a). Modes of rapid polygenic adaptation. Molecular Biology and Evolution, 34(12):3169–3175.

Jain, K. and Stephan, W. (2017b). Rapid Adaptation of a Polygenic Trait After a Sudden Environmental Shift. Genetics, 206(1):389–406.

Jakšić, A. M. and Schlötterer, C. (2016). The interplay of temperature and genotype on patterns of alternative splicing in *Drosophila melanogaster*. Genetics, 204(1):315–325.

Johansson, A. M., Pettersson, M. E., Siegel, P. B., and Carlborg, Ö. (2010). Genome-wide effects of long-term divergent selection. PLOS Genetics, 6(11):1–12.

Jónás, Á., Taus, T., Kosiol, C., Schlötterer, C., and Futschik, A. (2016). Estimating the effective population size from temporal allele frequency changes in experimental evolution. Genetics, 204(2):723–735.

Kawecki, T. J., Lenski, R. E., Ebert, D., Hollis, B., Olivieri, I., and Whitlock, M. C. (2012). Experimental evolution. Trends in Ecology & Evolution, 27(10):547 – 560.

Kelly, J. K. and Hughes, K. A. (2018). Pervasive linked selection and intermediate-frequency alleles are implicated in an evolve-and-resequencing experiment of *Drosophila simulans*. Genetics, 211(3):943–963.

Kessner, D. and Novembre, J. (2015). Power analysis of artificial selection experiments using efficient whole genome simulation of quantitative traits. Genetics, 199(4):991–1005.

Kofler, R., Pandey, R. V., and Schlötterer, C. (2011). PoPoolation2: identifying differentiation between populations using sequencing of pooled DNA samples (Pool-Seq). Bioinformatics (Oxford, England), 27(24):3435–3436.

Kofler, R. and Schlötterer, C. (2014). A guide for the design of evolve and resequencing studies. Molecular Biology and Evolution, 31(2):474–483.

Lang, G. I., Rice, D. P., Hickman, M. J., Sodergren, E., Weinstock, G. M., Botstein, D., and Desai, M. M. (2013). Pervasive genetic hitchhiking and clonal interference in forty evolving yeast populations. Nature, 500(7464):571–574.

Long, A., Liti, G., Luptak, A., and Tenaillon, O. (2015). Elucidating the molecular architecture of adaptation via evolve and resequence experiments. Nature Reviews Genetics, 16(10):567–582.

Martins, N. E., Faria, V. G., Nolte, V., Schlötterer, C., Teixeira, L., Sucena, É., and Magalhães, S. (2014). Host adaptation to viruses relies on few genes with different cross-resistance properties. Proceedings of the National Academy of Sciences, 111(16):5938–5943.

Michalak, P., Kang, L., Schou, M. F., Garner, H. R., and Loeschcke, V. (2019). Genomic signatures of experimental adaptive radiation in *Drosophila*. Molecular Ecology, 28(3):600–614.

Nuzhdin, S. V. and Turner, T. L. (2013). Promises and limitations of hitchhiking mapping. Current opinion in genetics & development, 23(6):694–699.

Orozco-Terwengel, P., Kapun, M., Nolte, V., Kofler, R., Flatt, T., and Schlötterer, C. (2012). Adaptation of *Drosophila* to a novel laboratory environment reveals temporally heterogeneous trajectories of selected alleles. Molecular Ecology, 21(20):4931–4941.

Papkou, A., Guzella, T., Yang, W., Koepper, S., Pees, B., Schalkowski, R., Barg, M.-C., Rosenstiel, P. C., Teotónio, H., and Schulenburg, H. (2019). The genomic basis of Red Queen dynamics during rapid reciprocal host–pathogen coevolution. Proceedings of the National Academy of Sciences, 116(3):923–928.

Pollard, K. S. and Laan, M. J. V. D. (2005). Analysis of Genomic Data with Applications in R. U.C. Berkeley Division of Biostatistics Working Paper Series, Working Paper 167.

Rêgo, A., Messina, F. J., and Gompert, Z. (2019). Dynamics of genomic change during evolutionary rescue in the seed beetle *Callosobruchus maculatus*. Molecular Ecology, 28(9):2136–2154.

Schlötterer, C., Kofler, R., Versace, E., Tobler, R., and Franssen, S. U. (2015). Combining experimental evolution with next-generation sequencing: a powerful tool to study adaptation from standing genetic variation. Heredity, 114(5):431–440.

Schlötterer, C., Tobler, R., Kofler, R., and Nolte, V. (2014). Sequencing pools of individuals — mining genome-wide polymorphism data without big funding. Nature Reviews Genetics, 15(11):749–763.

Seabra, S. G., Fragata, I., Antunes, M. A., Faria, G. S., Santos, M. A., Sousa, V. C., Simões, P., and Matos, M. (2019). Different Genomic Changes Underlie Adaptive Evolution in Populations of Contrasting History. Molecular Biology and Evolution, 36(6):1358–1358.

Taus, T., Futschik, A., and Schlötterer, C. (2017). Quantifying Selection with Pool-Seq Time Series Data. Molecular Biology and Evolution, 34(11):3023–3034.

Thornton, K. R. (2019). Polygenic adaptation to an environmental shift: Temporal dynamics of variation under Gaussian stabilizing selection and additive effects on a single trait. Genetics, 213(4):1513–1530.

Tibshirani, R., Walther, G., and Hastie, T. (2001). Estimating the number of clusters in a data set via the gap statistic. Journal of the Royal Statistical Society Series B, 63:411–423.

Tobler, R., Franssen, S. U., Nolte, V., and Schlötterer, C. (2014). Patterns of Linkage Disequilibrium and Long Range Hitchhiking in Evolving Experimental *Drosophila melanogaster* Populations. Molecular Biology and Evolution, 32(2):495–509.

Turner, T. L. and Miller, P. M. (2012). Investigating natural variation in *Drosophila* courtship song by the evolve and resequence approach. Genetics, 191(2):633–642.

Turner, T. L., Stewart, A. D., Fields, A. T., Rice, W. R., and Tarone, A. M. (2011). Population-based resequencing of experimentally evolved populations reveals the genetic basis of body size variation in *Drosophila melanogaster*. PLOS Genetics, 7(3):1–10.

Vlachos, C., Burny, C., Pelizzola, M., Borges, R., Futschik, A., Kofler, R., and Schlötterer, C. (2019). Benchmarking software tools for detecting and quantifying selection in evolve and resequencing studies. Genome Biology, 20(1):169.

Vlachos, C. and Kofler, R. (2018). Mimicree2: Genome-wide forward simulations of evolve and resequencing studies. PLOS Computational Biology, 14(8):1–10.

Vlachos, C. and Kofler, R. (2019). Optimizing the Power to Identify the Genetic Basis of Complex Traits with Evolve and Resequence Studies. Molecular biology and evolution, 36(12):2890– 2905.

