## Supplemental Information for "Low concordance of short-term and long-term selection responses in experimental *Drosophila* populations"

### Supplementary Material and Methods

#### Identification of candidate SNPs

SNPs were called applying the following criteria: base quality of 40 in at least one replicate, a read depth between the $2^{nd}$ and $98^{th}$ percentile, and the minor allele is supported by at least ten reads. Repeats, transposable elements, SNPs specific to Y-translocated genes (Tobler et al., 2017), and 5-bp regions around indels were excluded from the analysis to increase the robustness of the SNP set (for further details, see Barghi et al. (2019)), resulting in 5,096,200 SNPs on chromosomes X, 2, 3, and 4.

### References

Barghi, N., Tobler, R., Nolte, V., Jakšić, A. M., Mallard, F., Otte, K. A., Dolezal, M., Taus, T., Kofler, R., and Schlötterer, C. (2019). Genetic redundancy fuels polygenic adaptation in *Drosophila*. *PLOS Biology*, 17(2):1–31.

Tobler, R., Nolte, V., and Schlötterer, C. (2017). High rate of translocation-based gene birth on the *Drosophila* Y chromosome. *Proceedings of the National Academy of Sciences*, 114(44):11721–11726.

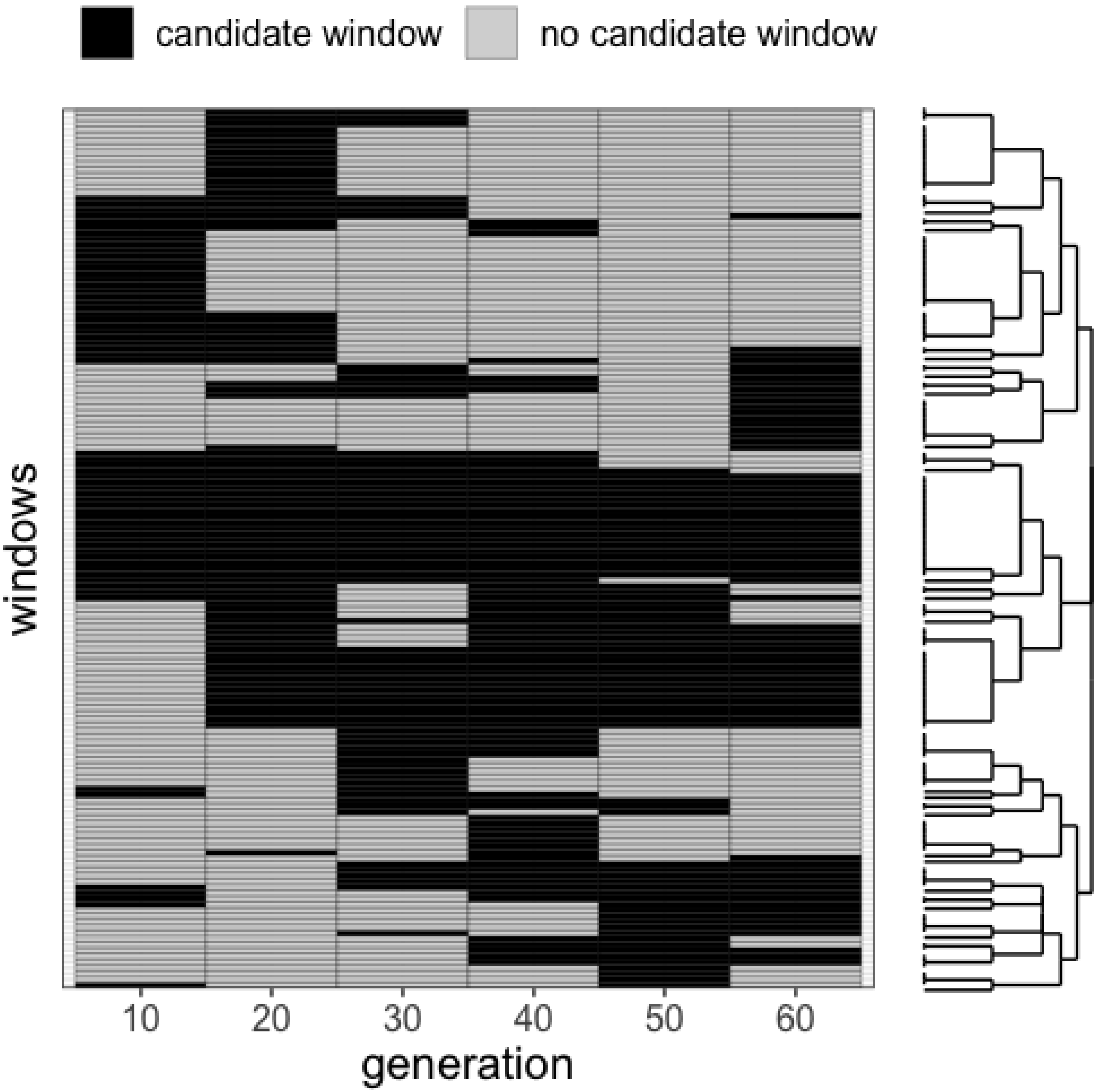

**Figure S1:** Time resolved representation of all windows that were classified as candidate window for at least one time point: each window is shown in a single row for generation 10 to 60. Windows that are enriched for candidate SNPs at a particular time point are colored black; not enriched windows are colored grey. Windows with similar candidate window patterns are grouped together.

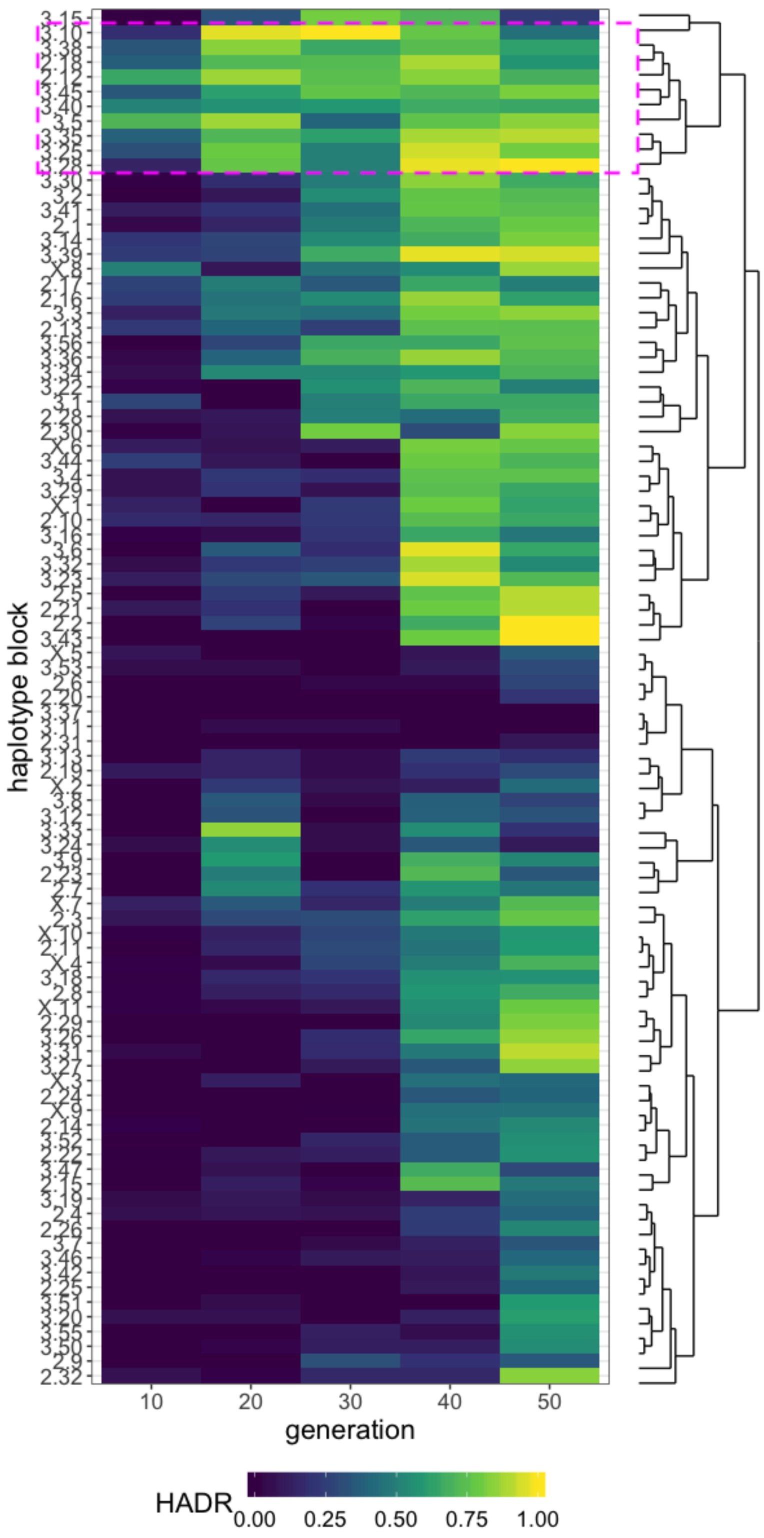

**Figure S2:** Empirical data: Haplotype discovery rate (HADR) from generation 10 to generation 50. The x-axis shows the five intermediate time points, the y-axis the haplotype blocks determined by Barghi et al. (2019). Haplotype blocks are grouped by similar HADR dynamics over time. Tiles are colored by HADR score. The 10 haplotype blocks of the Early Detected Haplotypes (EDHA) cluster (n=10) are highlighted by a pink frame.

While the HADR of most haplotype blocks increases with time, for nine haplotype blocks the HADR score fluctuated (2.7, 2.15, 2.23, 3.8, 3.9, 3.12, 3.24, 3.33, and 3.47). Because these haplotype blocks are mainly based on marginally significant candidate SNPs (Table S8), they are more sensitive to random fluctuations, partly due to experimental noise, partly due to stochasticity of the simulations used to determine the significance threshold.

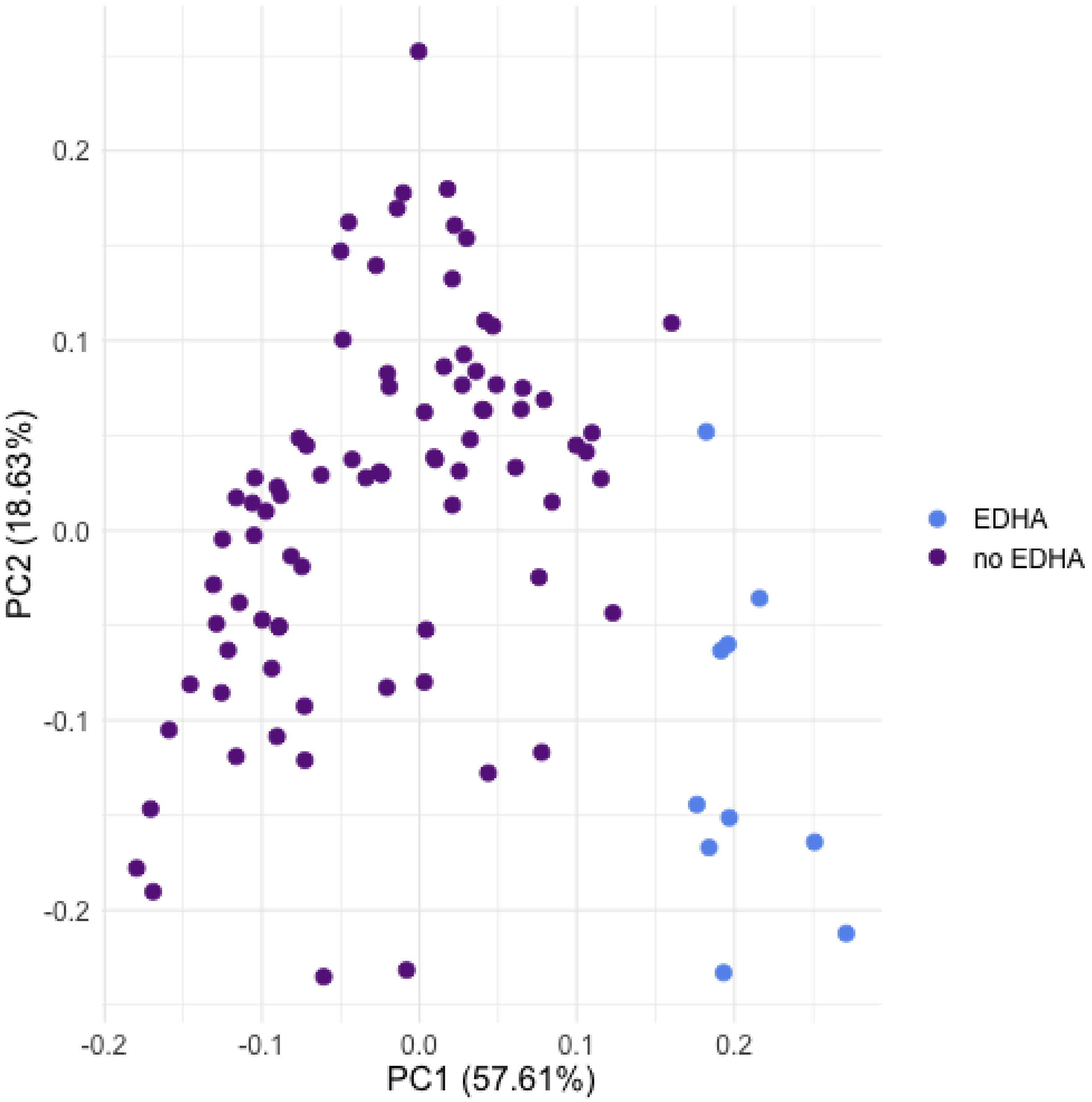

**Figure S3:** The first two principle components from a PCA based on the haplotype discovery rate (HADR) from generation 10 to generation 50. Each haplotype block is represented by a single dot. Blocks of the early detectable haplotype block (EDHA) cluster are colored in blue, others in purple.

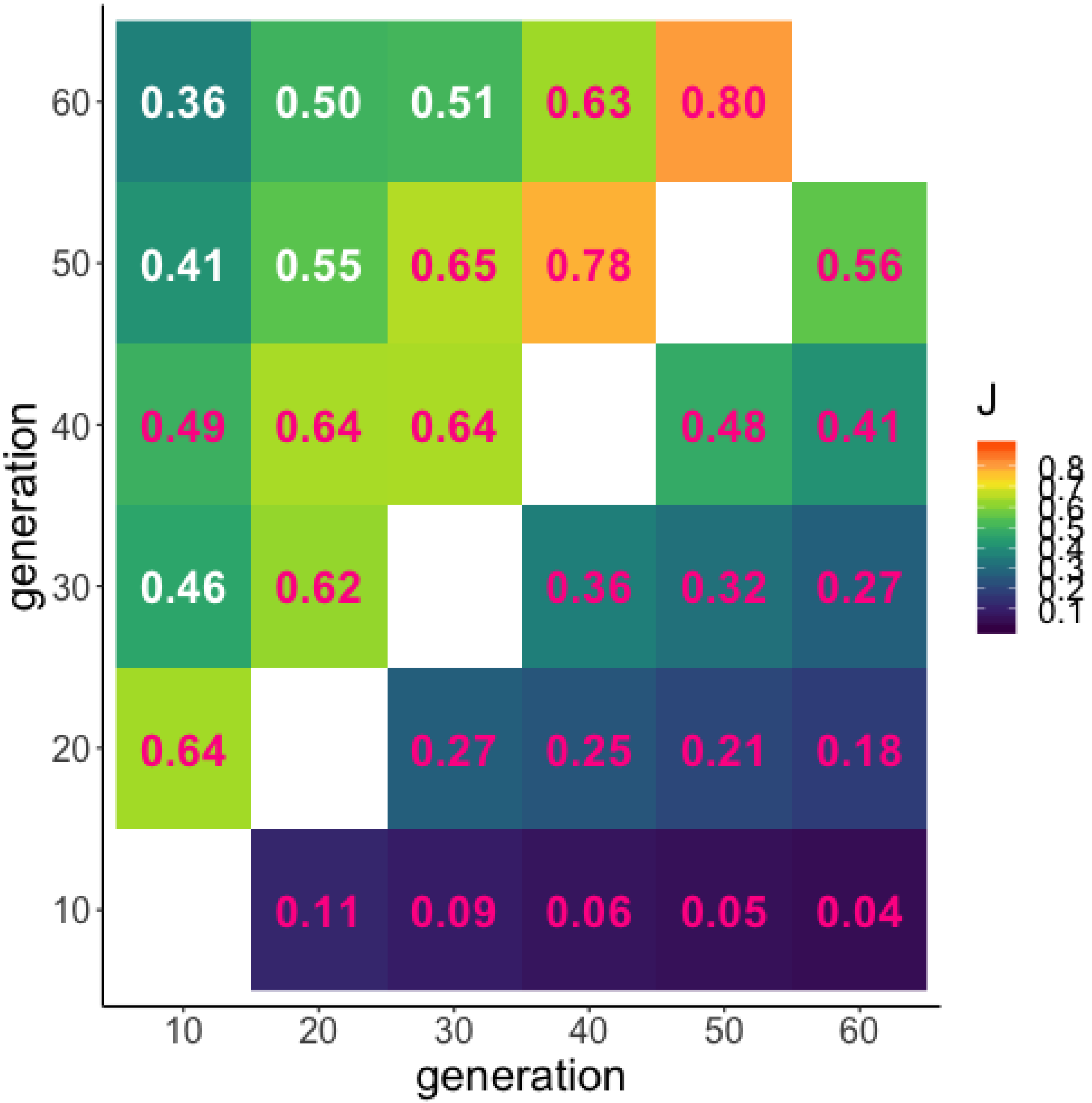

**Figure S4:** Jaccard index (J) for pairwise comparisons of candidate sets based on a simulated Evolve and Resequence experiment (filtered for SNPs segregating in all generations and time points). The top triangle shows candidate window sets, the bottom triangle candidate SNP sets. Significant similarities (*p*<0.05 after multiple testing correction, 10 000 bootstraps) are marked in pink.

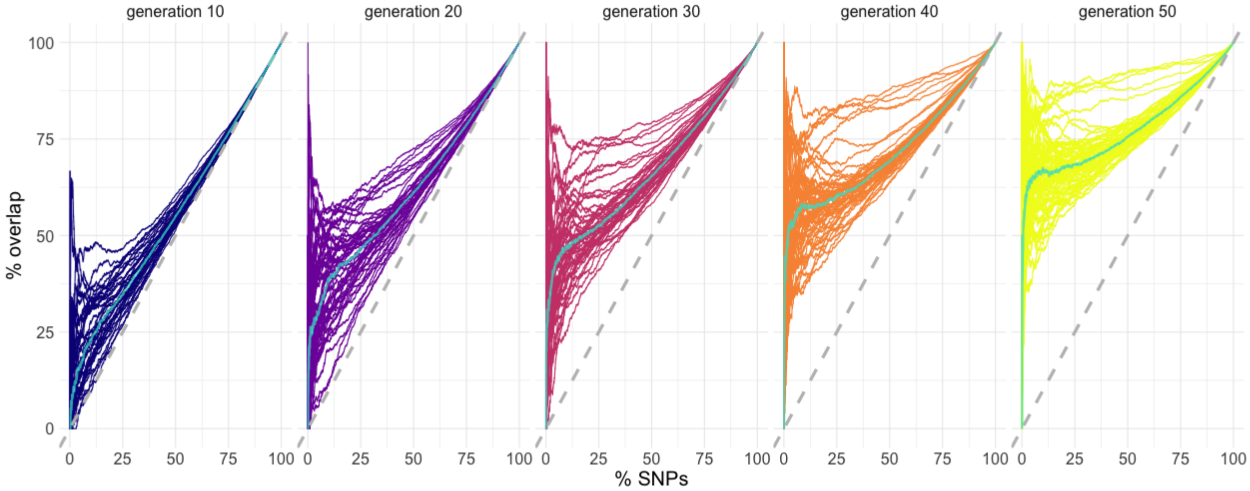

**Figure S5:** The rank of candidate SNPs becomes more congruent with time in the simulated data. In this ROC-like graph, the ranking of all candidate SNPs in candidate windows is compared. Each panel shows one intermediate time point compared to generation 60. The overlap (in percent) for each candidate window is indicated by a separate line. The median overlap (turquoise line) monotonically increases with experimental duration, demonstrating that the ranking of candidate SNPs is more robust for advanced generations. The grey, dashed lines shows the expected overlap in SNP ranking if every variant at generation 60 is recapitulated in a previous time point.

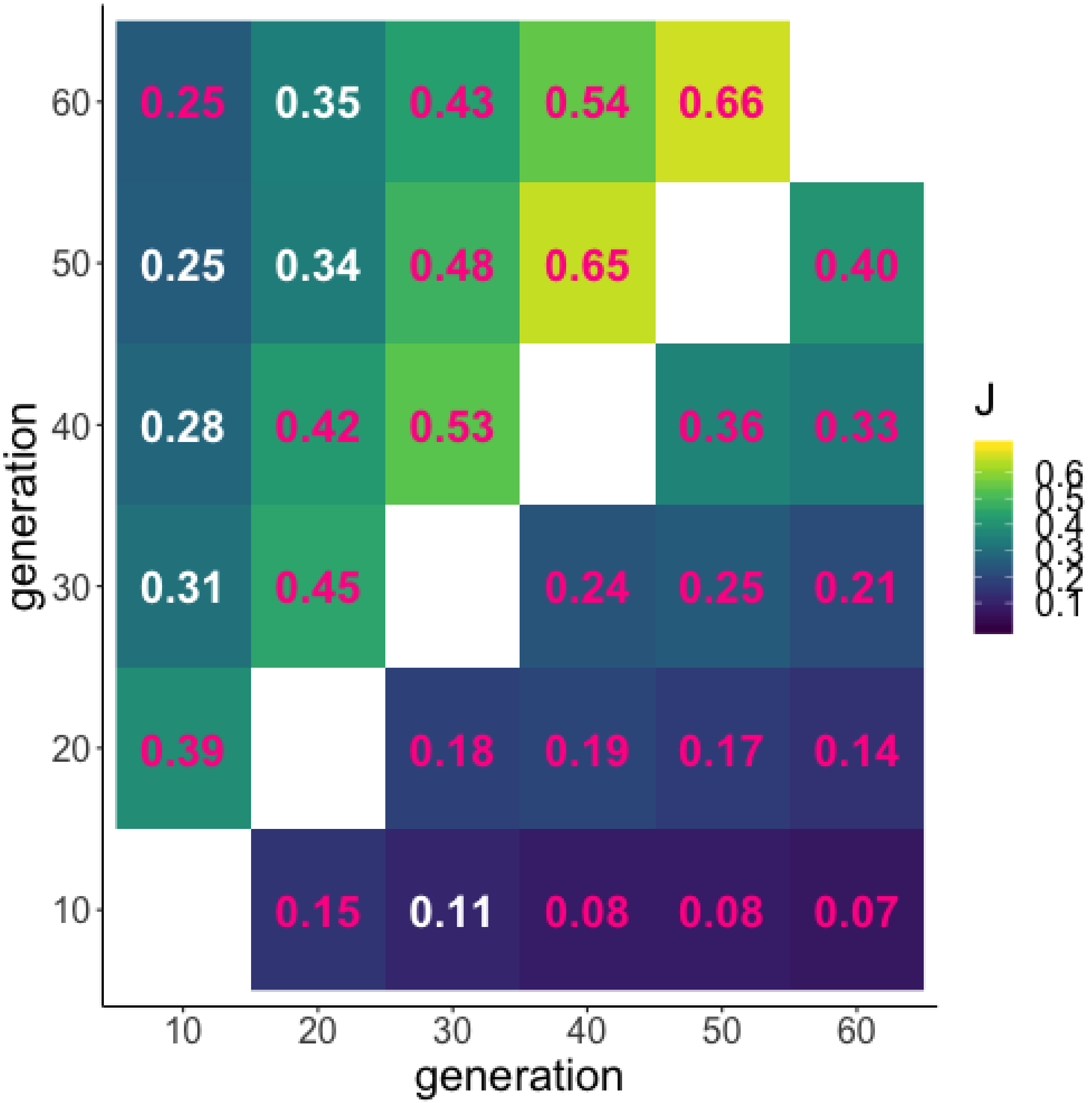

**Figure S6:** Jaccard index (J) for pairwise comparisons of empirical candidate sets (not filtered for SNPs segregating in all generations and time points). The top triangle shows candidate window sets, the bottom triangle candidate SNP sets. Significant similarities ($p$<0.05 after multiple testing correction, 10 000 bootstraps) are marked in pink.

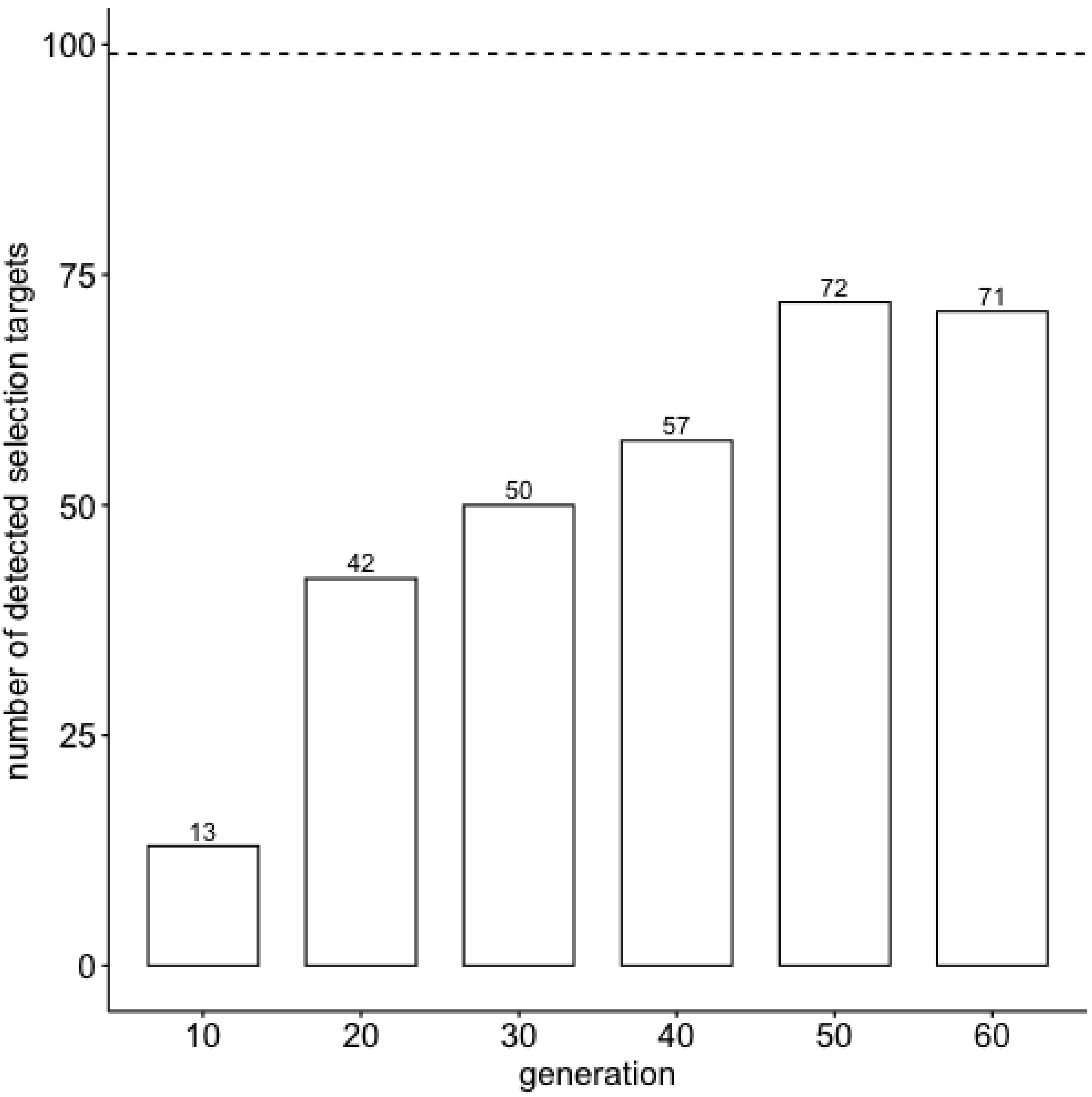

**Figure S7:** Number of detected causative SNPs for generation 10 to 60. 99 causative SNPs were simulated.

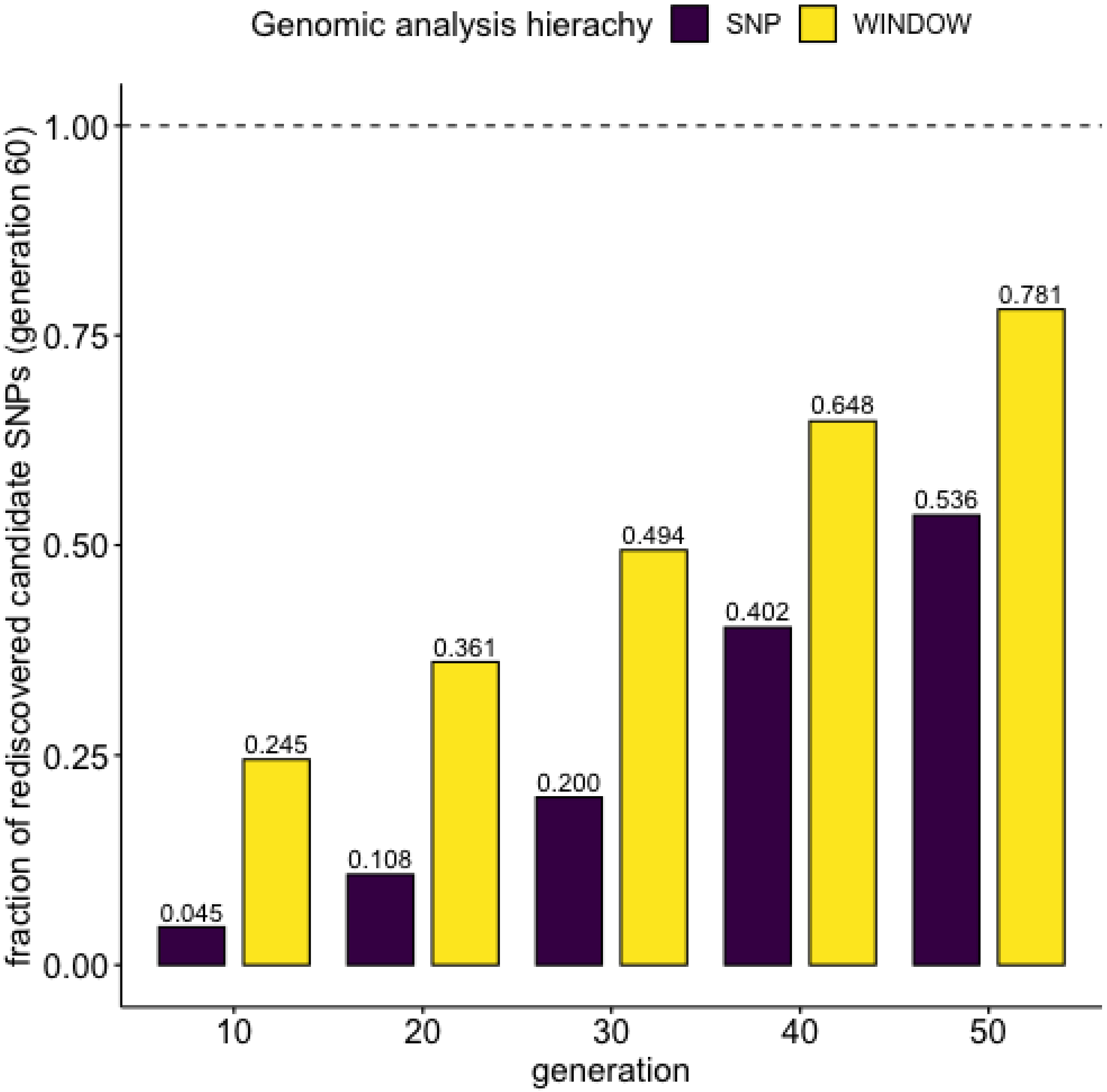

**Figure S8:** Less than 5% of candidate SNPs in generation 60 are detected consistently at every generation. The bars depict the ratio of candidates at generation 60 (SNPs in purple, windows in yellow) that consistently remain candidates at consecutive earlier time points (e.g. 40.2 % of the candidate SNPs at generation 60 are also candidates in generation 50 and 40). Window-based analyses are more consistent than SNP-based analyses.

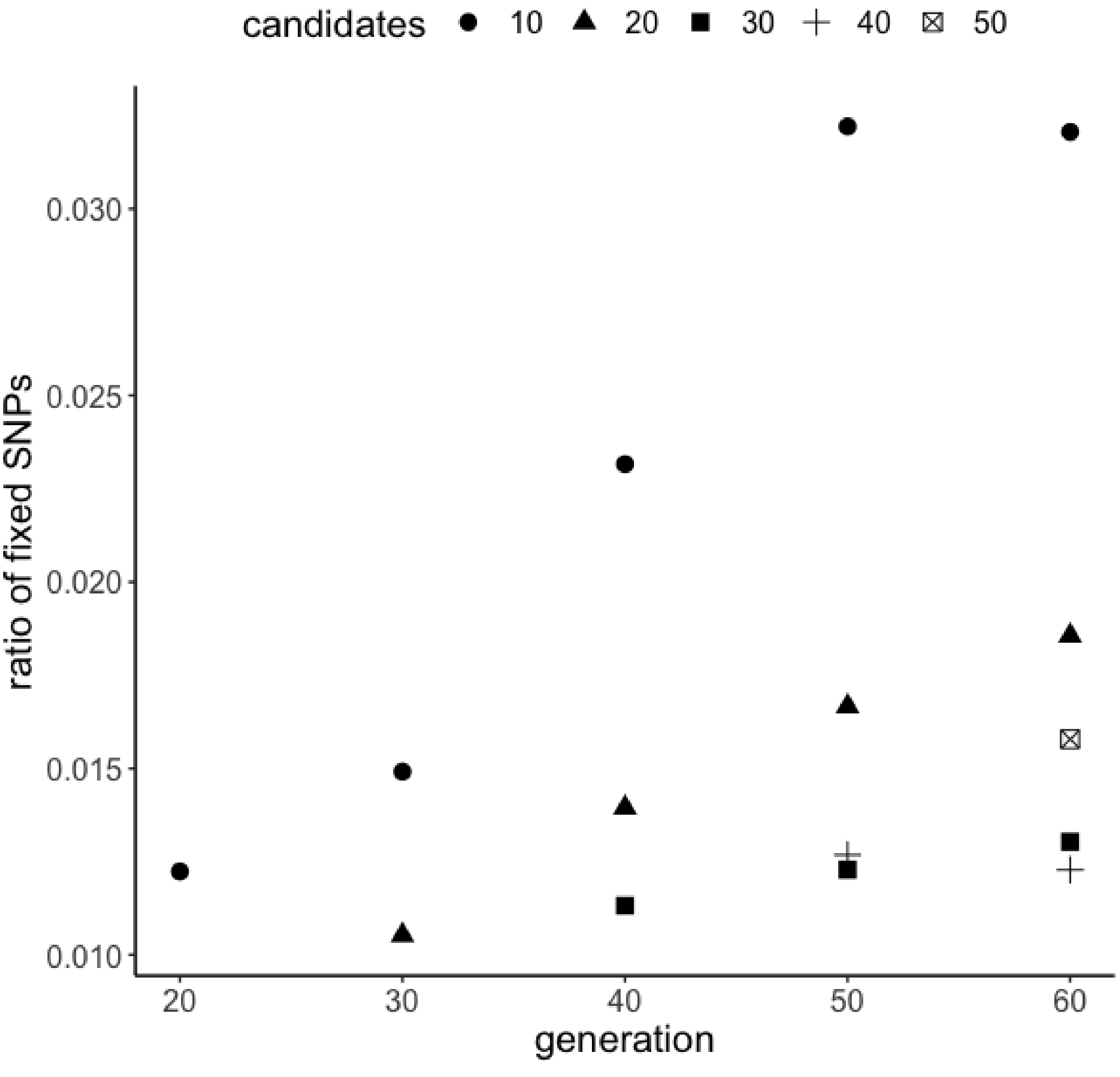

**Figure S9:** Less than 3.5 % of candidate SNPs in generation 10 are fixed at later generations. The points show the ratio of candidate SNPs that are fixed (frequency ≥ 0.99) in at least 8 out of 10 replicates at later time points.

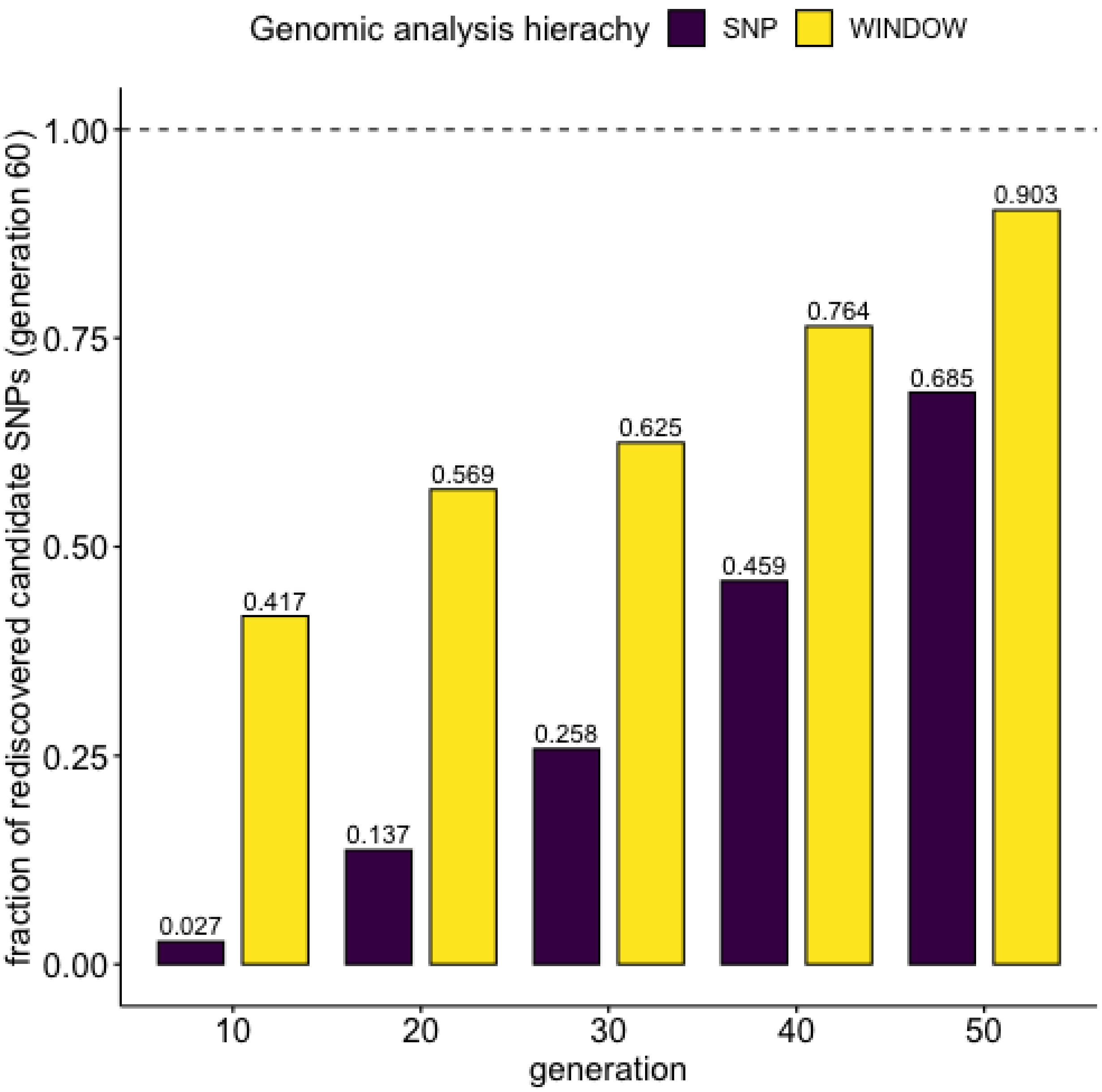

**Figure S10:** Less than 3% of candidate SNPs in generation 60 are detected consistently at every generation in our simulated Evolve and Resequence experiment. The bars depict the ratio of candidates at generation 60 (SNPs in purple, windows in yellow) that consistently remain candidates at consecutive earlier time points (e.g. 45.9 % of generation 60 candidate SNPs are also candidates in generation 50 and 40). Window-based analyses are more consistent than SNP-based analyses.

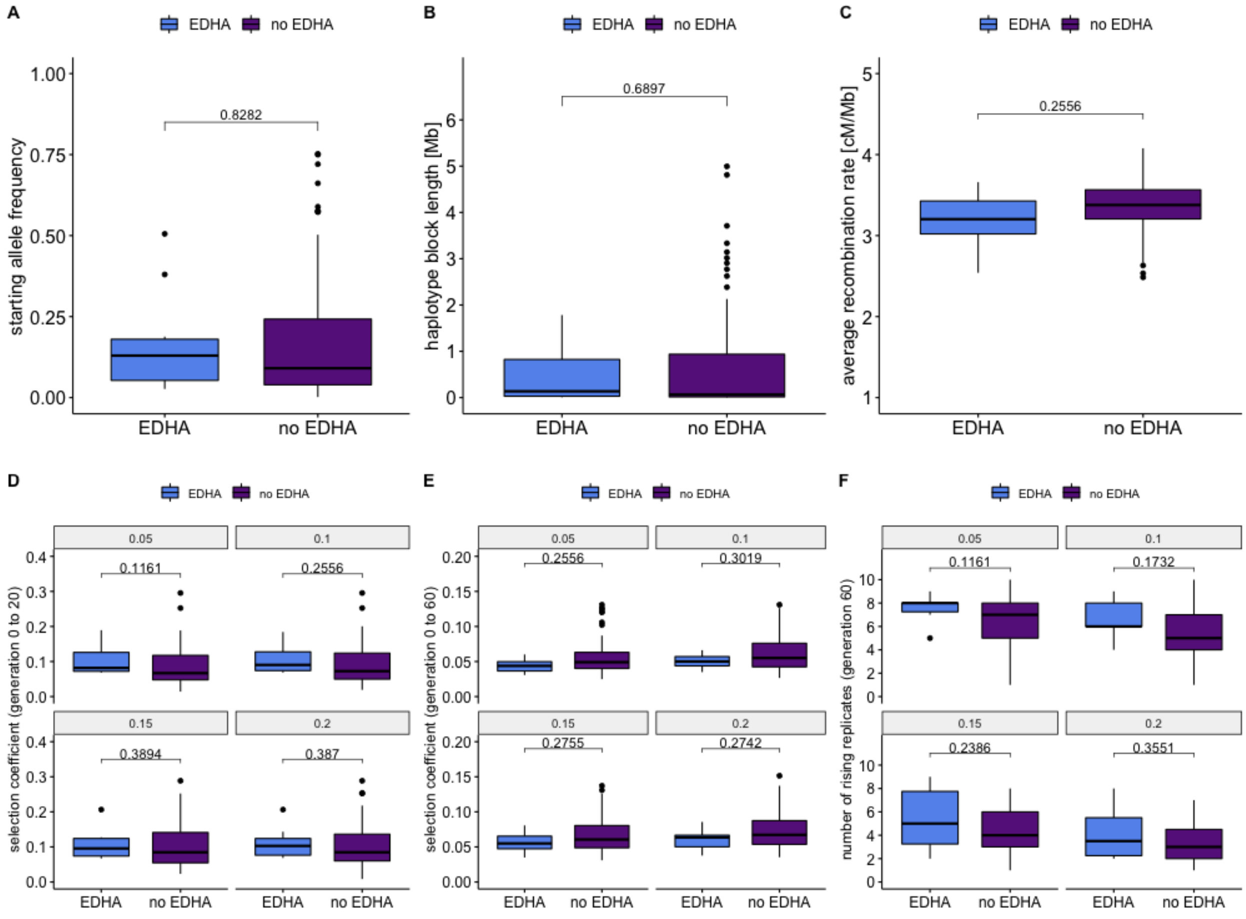

**Figure S11:** Early detectable haplotype blocks (EDHAs) do not differ in starting allele frequency (**A**), block length (**B**), average recombination rate (**C**), strength of selection (both early (**D**) and late (**E**) in the experiment), or number of rising replicates in generation 60 (**F**) from other haplotype blocks. Values above the boxplots represent the two-tailed Mann-Whitney test *p*-values corrected for multiple testing with the Benjamini-Hochberg procedure.

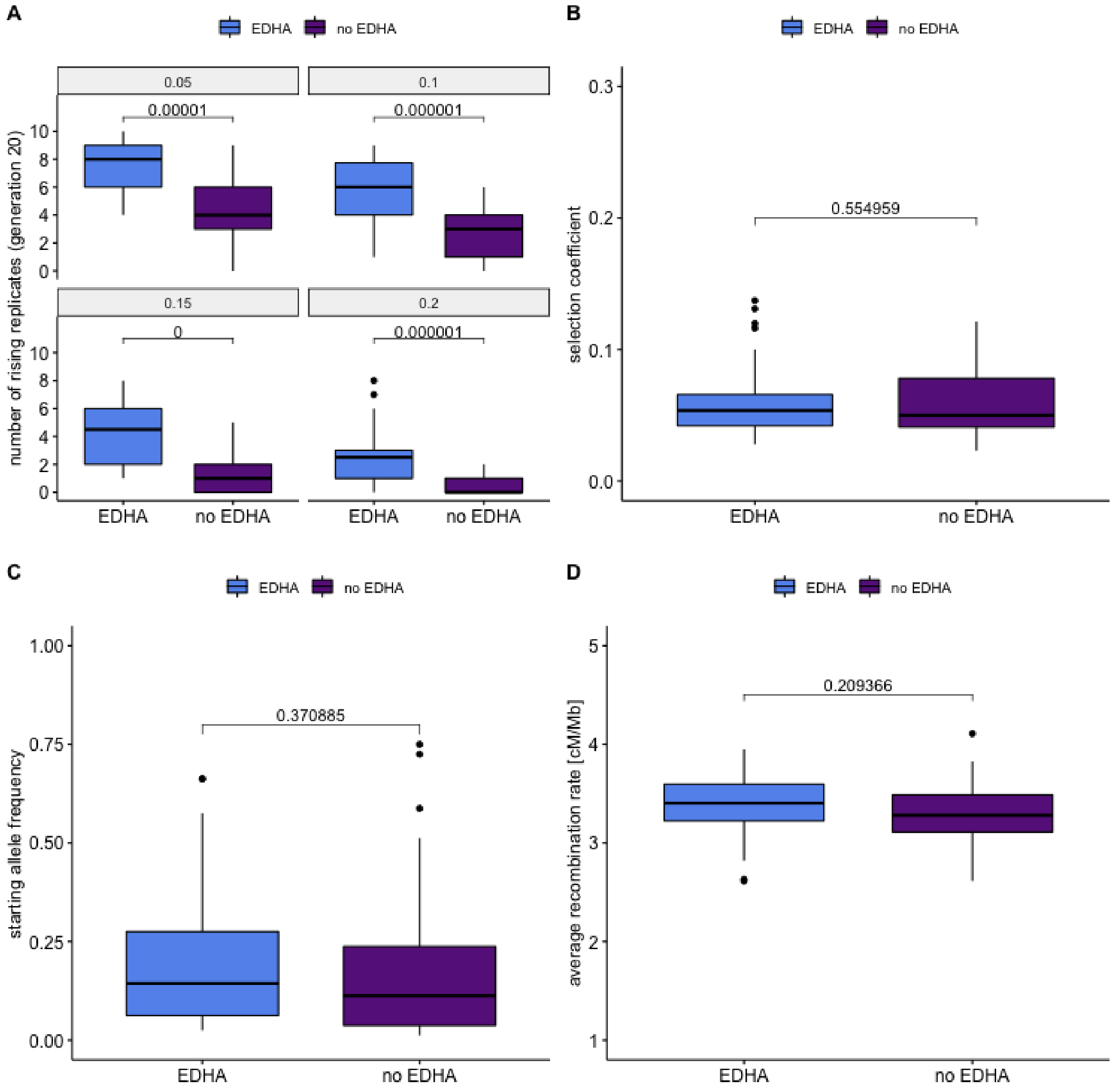

**Figure S12:** After 20 generations, early detectable targets respond in more replicates than other targets in the simulated Evolve and Resequence experiment – regardless of the chosen allele frequency change cutoff (**A**). Early detectable targets do not differ in selection strength (**B**), starting allele frequency (**C**) or average recombination rate (**D**) from other selected targets. Values above the boxplots represent the two-tailed Mann-Whitney test $p$-values corrected for multiple testing with the Benjamini-Hochberg procedure.

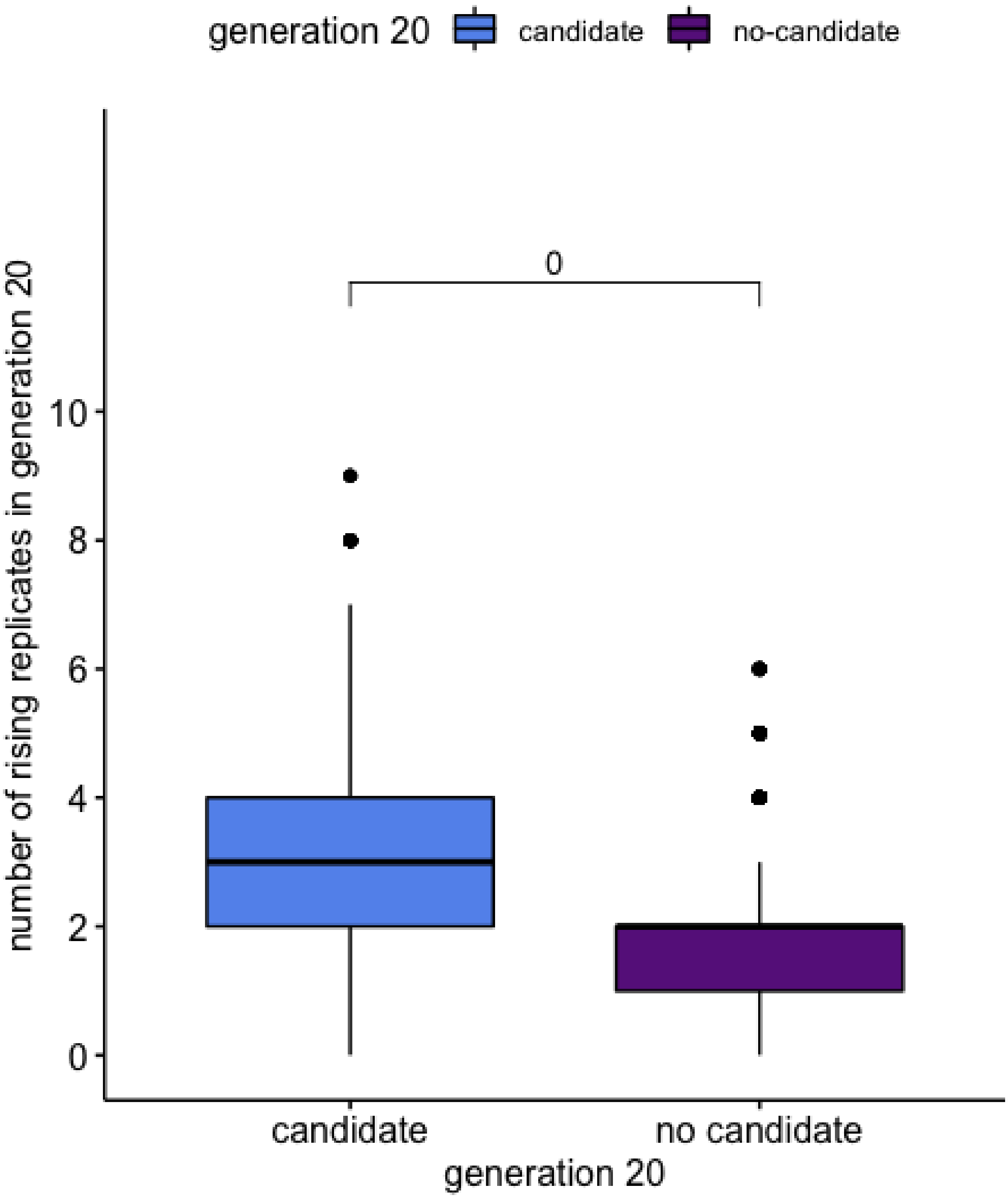

**Figure S13:** After 20 generations, detectable loci (candidates, FDR≤5%) respond in more replicates (allele frequency change of at least 5%) than not detectable loci, despite having the same starting allele frequency (10%), and selection coefficient (0.05) in computer simulations (100 000 loci simulated). The p-value above the boxplots represents the two-tailed Mann-Whitney test.

**Table S1:** $N_e$ estimates for the X-chromosome and autosomes averaged across 10 replicates at different time points

| generation | X | autosome |
|---|---|---|
| 10 | 257 | 278 |
| 20 | 230 | 261 |
| 30 | 237 | 266 |
| 40 | 239 | 260 |
| 50 | 245 | 269 |
| 60 | 262 | 291 |

**Table S2:** Replicate-specific autosomal $N_e$ estimates for different time points

| generation | R1 | R2 | R3 | R4 | R5 | R6 | R7 | R8 | R9 | R10 |
|---|---|---|---|---|---|---|---|---|---|---|
| 10 | 334 | 431 | 333 | 279 | 213 | 251 | 236 | 257 | 262 | 188 |
| 20 | 309 | 310 | 240 | 208 | 226 | 274 | 221 | 274 | 316 | 235 |
| 30 | 337 | 304 | 238 | 210 | 234 | 290 | 239 | 243 | 288 | 275 |
| 40 | 341 | 306 | 238 | 205 | 227 | 287 | 255 | 211 | 270 | 257 |
| 50 | 356 | 306 | 237 | 235 | 215 | 289 | 281 | 226 | 288 | 257 |
| 60 | 381 | 336 | 247 | 255 | 246 | 307 | 297 | 244 | 310 | 287 |

**Table S3:** Replicate-specific X $N_e$ estimates for different time points

| generation | R1 | R2 | R3 | R4 | R5 | R6 | R7 | R8 | R9 | R10 |
|---|---|---|---|---|---|---|---|---|---|---|
| 10 | 291 | 411 | 279 | 261 | 209 | 228 | 224 | 251 | 211 | 201 |
| 20 | 181 | 269 | 220 | 226 | 236 | 191 | 255 | 253 | 240 | 230 |
| 30 | 190 | 281 | 251 | 207 | 237 | 218 | 233 | 239 | 266 | 253 |
| 40 | 191 | 268 | 265 | 213 | 220 | 230 | 252 | 217 | 286 | 248 |
| 50 | 203 | 279 | 260 | 231 | 214 | 236 | 256 | 229 | 292 | 250 |
| 60 | 217 | 280 | 269 | 240 | 208 | 274 | 270 | 241 | 358 | 268 |

**Table S4:** Fraction of candidate windows at generation 60, that are also candidate windows in all previous generations. a: windows size = 5 000 SNPs. SNPs do not necessarily segregate in all replicates and time points. b: window size =13 025 SNPs (a window with 13 025 SNPs result on average in a window length of 250 kb: that is half the step size in the sliding window approach used by Barghi et al. (2019) to reconstruct selected haplotype blocks) . SNPs do not necessarily segregate in all replicates and time points. c: window size =5 000 SNPs, which segregate in all replicates and time points.

| a | b | c |
|---|---|---|
| 0.24% | 0.28% | 0.24% |

**Table S5:** Absolute number and fraction of candidate windows at each generation. a: windows size = 5 000 SNPs. SNPs do not necessarily segregate in all replicates and time points. b: window size =13 025 SNPs. SNPs do not necessarily segregate in all replicates and time points. c: window size =5 000 SNPs, which segregate in all replicates and time points.

| generation | a | b | c |
|---|---|---|---|
| 10 | 186 (0.18) | 78 (0.20) | 62 (0.17) |
| 20 | 224 (0.22) | 109 (0.28) | 83 (0.23) |
| 30 | 196 (0.19) | 90 (0.23) | 72 (0.20) |
| 40 | 235 (0.23) | 111 (0.28) | 83 (0.23) |
| 50 | 225 (0.22) | 92 (0.23) | 69 (0.19) |
| 60 | 233 (0.23) | 97 (0.25) | 74 (0.21) |

**Table S6:** Number of candidate SNPs at each time point

| generation | number of candidate SNPs |
|---|---|
| 10 | 27 290 |
| 20 | 42 736 |
| 30 | 31 002 |
| 40 | 64 974 |
| 50 | 48 344 |
| 60 | 56 166 |

**Table S7:** Number of candidate SNPs at each time point in the simulated Evolve and Resequence data.
true positives: SNPs that are candidate SNPs and s>0
false positives: SNPs that are candidate SNPs and s=0

| generation | number of candidates SNPs | true positives | false positives |
|---|---|---|---|
| 10 | 35 954 | 13 | 35 941 |
| 20 | 108 129 | 42 | 108 087 |
| 30 | 106 441 | 50 | 106 391 |
| 40 | 142 696 | 57 | 142 639 |
| 50 | 177 052 | 72 | 176 980 |
| 60 | 194 573 | 71 | 194 502 |

**Table S8:** Highest -$\log_{10}(p\text{–value})$ in haplotype blocks with HADR fluctuation over time. The first row in the table shows the applied 5 % FDR cut-off for autosomes. The second row contains the highest -$\log_{10}(p\text{–value})$ observed. Row three to eleven depict the highest -$\log_{10}(p\text{–values})$ for different haplotype blocks. We only depict haplotype blocks with pronounced HADR fluctuations at certain time points.

| generation | 20 | 30 | 40 | 50 | 60 |
|---|---|---|---|---|---|
| cut-off | 10 | 14 | 15 | 18 | 19 |
| max | 74.06 | 66.74 | 78 | 102.5 | 112.85 |
| 3.8 | 15.95 | 14.1 | 23.37 | | |
| 3.12 | 12.09 | 13.09 | 22.02 | | |
| 3.33 | 18.65 | 14.69 | 19.34 | | |
| 3.24 | 22.86 | 12.48 | 20.87 | | |
| 3.9 | 15.30 | 13.58 | 24.56 | | |
| 2.23 | 19.17 | 13.55 | 24.74 | | |
| 2.7 | 17.05 | 18.76 | 23.00 | | |
| 3.47 | | | 21.44 | 26.8 | 30.85 |
| 2.15 | | | 32.21 | 32.63 | 30.03 |
